# The population tracking model: A simple, scalable statistical model for neural population data

**DOI:** 10.1101/064717

**Authors:** Cian O’Donnell, J. Tiago Gonçalves, Nick Whiteley, Carlos Portera-Cailliau, Terrence J. Sejnowski

**Affiliations:** Department of Computer Science, Bristol, UK; School of Mathematics, University of Bristol, Bristol, UK; Howard Hughes Medical Institute, La Jolla CA, USA; Salk Institute for Biological Studies, La Jolla CA, USA; Departments of Neurology and Neurobiology, David Geffen School of Medicine at UCLA, Los Angeles, CA, USA; Division of Biological Sciences, University of California at San Diego, La Jolla, CA, USA

## Abstract

Our understanding of neural population coding has been limited by a lack of analysis methods to characterize spiking data from large populations. The biggest challenge comes from the fact that the number of possible network activity patterns scales exponentially with the number of neurons recorded (∼ 2^Neurons^). Here we introduce a new statistical method for characterizing neural population activity that requires semi-independent fitting of only as many parameters as the square of the number of neurons, so requiring drastically smaller data sets and minimal computation time. The model works by matching the population rate (the number of neurons synchronously active) and the probability that each individual neuron fires given the population rate. We found that this model can accurately fit synthetic data from up to 1000 neurons. We also found that the model could rapidly decode visual stimuli from neural population data from macaque primary visual cortex, ∼ 65 ms after stimulus onset. Finally, we used the model to estimate the entropy of neural population activity in developing mouse somatosensory cortex and surprisingly found that it first increases, then decreases during development. This statistical model opens new options for interrogating neural population data, and can bolster the use of modern large-scale in vivo Ca^2+^ and voltage imaging tools.

## 1 Introduction

Brains encode and process information as electrical activity over populations of their neurons (Churchland and Sejnowski, 1994; Averbeck et al., 2006). Although understanding the structure of this neural code has long been a central goal of neuroscience, historical progress has been impeded by limitations in recording techniques. Traditional extracellular recording electrodes allowed isolation of only one or a few neurons at a time (Stevenson and Kording, 2011). Given that the human brain has on the order of 10^11^ neurons, the contribution of such small groups of neurons to brain processing is likely minimal. To get a more complete picture we would instead like to simultaneously observe the activity of large populations of neurons. Although the ideal scenario — recording every neuron in the brain — is out of reach for now, recent developments in both electrical and optical recording technologies have increased the typical size of population recording so that many laboratories now routinely record from hundreds or even thousands of neurons (Stevenson and Kording, 2011). The advent of these big neural data has introduced a new problem: how to analyze them.

The most commonly applied analysis to neural population data is to simply examine the activity properties of each neuron in turn, as if they were recorded in separate animals. However responses of nearby neurons to sensory stimuli are often significantly correlated, implying that neurons do not process information independently (Perkel et al., 1967; Gerstein and Perkel, 1969, 1972; Singer, 1999; Cohen and Kohn, 2011). As a result, performing a cell-by-cell analysis amounts to throwing away potentially valuable information on the collective behavior of the recorded neurons. These correlations are important because they put strong functional constraints on neural coding (Zohary et al., 1994; Averbeck et al., 2006).

If we consider each neuron to have two spiking activity states, ON or OFF, then a population of N neurons as a whole can have *2^N^* possible ON/OFF patterns at any moment in time. The probability of seeing any particular one of these population activity patterns depends on the brain circuit examined, the stimuli the animal is subject to, and perhaps also the internal brain state of the animal. Neural correlations and sparse firing imply that the probability of some activity patterns are more likely than others. To help understand the neural code we would like to be able to estimate the probability distribution across all 2^*N*^ patterns, *P_true_*. For small *N*, the probability of each pattern can be estimated by simply counting each time it appears, then dividing by the total number of timepoints recorded. However, since the number of possible patterns increases exponentially with *N*, this histogram method is experimentally intractable for populations larger than ∼ 10 neurons. For example, 20 neurons would require fitting 2^20^ ≈ 10^6^ parameters, one for each possible activity pattern. To accurately fit this model by counting patterns alone would require data recorded for many weeks or months. The problem gets worse for larger numbers of neurons: each additional neuron recorded requires a doubling in the recording time to reach the same level of statistical accuracy. This explosive scaling implies that we can never know the true distribution of pattern probabilities for a large number of neurons in a real brain.

This problem remained intractable until a seminal paper in 2006 demonstrated a possible solution: to fit a statistical model to the data that matches only some of the key low-order statistics, such as firing rates and pairwise correlations, and assume nothing else (Schneidman et al., 2006). The hope was that these basic statistics are sufficient for the model to capture the majority of structure present in the real data so that *P_moddel_* ≈ *P_true_*. Indeed early studies showed that such *pairwise maximum entropy* models could accurately capture activity pattern probabilities from recordings of 10–15 neurons in retina and cortex (Schneidman et al., 2006; Shlens et al., 2006; Tang et al., 2008; Yu et al., 2008). Unfortunately however, later studies found that performance of these pairwise models was poor for larger populations and in different activity regimes (Ohiorhenuan et al., 2010; Ganmor et al., 2011; Yu et al., 2011; Yeh et al., 2010), as predicted by theoretical work (Roudi et al., 2009; Macke et al., 2011a). As a consequence, variants of the pairwise maximum entropy models have been proposed that include higher-order correlation terms (Ganmor et al., 2011; Tkacik et al., 2013, 2014), but these are difficult to fit for large *N* and are not readily normalizable. Alternative approaches have also been developed that appear to provide better matches to data (Amari et al., 2003; Pillow et al., 2008; Macke et al., 2009, 2011b; Köster et al., 2014; Okun et al., 2012; Park et al., 2013; Okun et al., 2015; Schölvinck et al., 2015; Cui et al., 2016), but these suffer from similar shortcomings (Table 1). We suggest the following criteria for an ideal statistical model for neural population data:

1. It should accurately capture the structure in real neural population data.
2. Its fitting procedure should scale well to large N, meaning that the model’s parameters can be fit to data from large neural populations with a reasonable amount of data and computational resources.
3. Quantitative predictions can be made from the model after it is fit.

**Tab. 1:** Comparison of properties of various statistical models of neural activity. For the “Number of parameters” column’, *N* indicates the number neurons considered, ∼ indicates “scales with”, *D* indicates the number of coefficients per interaction term, and *R* indicates the number of timepoints across which temporal correlations are considered.

| Model | References | Number of parameters | Sampling possible? | Fit for large $N$? | Direct estimates of pattern probabilities? | Low-dimensional model of entire distribution? |
| --- | --- | --- | --- | --- | --- | --- |
| Pairwise maximum entropy | Schneidman et al. (2006); Shlens et al. (2006) | $\sim N^2$ | Yes | Difficult | Difficult | No |
| K-pairwise maximum entropy | Tkacik et al. (2013, 2014) | $\sim N^2$ | Yes | Difficult | Difficult | No |
| Spatiotemporal maximum entropy | Marre et al. (2009); Nasser et al. (2013) | $\sim RN^2$ | Yes | Difficult | Difficult | No |
| semi-Restricted Boltzmann Machine | Köster et al. (2014) | $\sim N^2$ | Yes | Difficult | Difficult | No |
| Reliable interaction model | Ganmor et al. (2011) | Data-dependent | No | Yes | Approximate | No |
| Generalized Linear Models | Pillow et al. (2008) | $\sim DN^2$ | Yes | Difficult | No | No |
| Dichotomized Gaussian | Amari et al. (2003); Macke et al. (2009) | $\sim N^2$ | Yes | Yes | No | No |
| Cascaded Logistic | Park et al. (2013) | $\sim N^2$ | Yes | Yes | Yes | No |
| Population coupling | Okun et al. (2012, 2015) | $3N$ | Yes | Yes | No | No |
| Population tracking | This study | $N^2$ | Yes | Yes | Yes | Yes |

No existing model meets all three of these demands (Table 1). Here we propose a novel, simple statistical method that does: the population tracking model. The model is specified by only *N*^2^ parameters: *N* to specify the distribution of number of neurons synchronously active, and a further *N*^2^ — *N* for the conditional probabilities that each individual neuron is ON given the population rate. Although no model with *N*^2^ parameters can ever fully capture all 2^*N*^ pattern probabilities, we find that the population tracking model strikes a good balance between accuracy, tractability, and usefulness: by design it matches key features of the data, its parameters can be easily fit for large *N*, it is normalizable allowing expression of pattern probabilities in closed form, and most surprisingly it allows estimation of measures of the entire probability distribution, as we demonstrate for neural populations as large as *N* = 1000.

The results sections of this paper is structured as follows. In section 2.1 we introduce the basic mathematical form of the model, and fit it to spiking data from macaque visual cortex as an illustration. In sections 2.2 and 2.3 we cover how the model parameters can be estimated from data, and how to sample synthetic data from the fitted model. In section 2.4 we show how a reduced 3*N*-parameter model of the entire 2^*N*^-dimensional pattern probability distribution can be derived from the model parameters, and how this reduced model can be used to estimate the population entropy, and the divergence between the model fits to two different datasets. In sections 2.5, 2.6 and 2.7 we show how the model’s estimates for entropy and pattern probabilities converge as a function of neuron number and time samples available. Finally, in sections 2.7 and 2.8 we show how the method can help give novel biological insights by applying it to two data sets: first we use the model to decode stimuli from the recorded electrophysiological spiking responses in macaque V1, and second, we analyze *in vivo* two-photon Ca^2+^ imaging data from mouse somatosensory cortex to explore how the entropy of neural population activity changes during development.

## 2 Results

### 2.1 Overview of the statistical model with example application to data

We consider parallel recordings of the electrical activity of a population of *N* neurons. If the recordings are made using electrophysiology, then spike sorting methods can be used to extract the times of action potentials emitted by each neuron from the raw voltage waveforms (Quiroga, 2012). If the data are recorded using imaging methods, for example via a Ca^2+^-sensitive fluorophore, then electrical spike times or neural firing rates can often be approximately inferred (Pnevmatikakis et al., 2016; Rahmati et al., 2016). Regardless of the way the in which the data are collected, at any particular timepoint in the recording some subset of these neurons may be active (ON), and the rest inactive (OFF). In the case of electrophysiologically recorded spike trains, the neurons considered ON might be those that emitted one or more spikes within a particular time bin Δ*t*. For fluorescence imaging data, a suitable threshold in the Δ*F*(*t*)/*F_0_* signal may be chosen to split neurons into ON and OFF groups, perhaps after also binning the data in time. Once we have binarized the neural activity data in this way, each neuron’s activity across time is reduced to a binary sequence of zeros and ones, where a zero represents silence and a one represents activity. For example, the ith neuron’s activity in the population might be x_*i*_ = 0,1,0, 0, 0,1,1, 0,1…. The length of the sequence *T* is simply the total number of time bins recorded. The brain might encode sensory information about the world in these patterns of neural population activity.

Next we can next group the neural population data into a large *N* × *T* matrix ***M*** where each row from *i* = 1 : *N* corresponds to a different neuron and each column from *j* = 1 : *T* corresponds to a different time point. At any particular time point (the *j*th column of ***M***), we could in principle see any possible pattern of inactive and active neurons, written as a vector of zeros and ones {*x*}_*j*_ = [*x_1j_*, *x*_2*j*_… *x_N_j*]^*T*^. In general, there will be *2^N^* possible patterns of population activity, or combinations of zeros and ones. In any given experiment, each particular pattern must have some ground-truth probability of appearing *P_true_*({*x*}), depending on the stimulus, animal’s brain state, and so on. We would like to estimate this 2^*N*^-dimensional probability distribution. However, since direct estimation is impossible, we instead fit the parameters of a simpler statistical model that implicitly specifies a different probability distribution over the patterns, *P_model_*({*x*}). The hope is that for typical neural data *P_model_*({*x*} ≈ *P_true_*({*x*}). In figure 1 we schematize the procedure for building and using such a model.

**Fig. 1:**
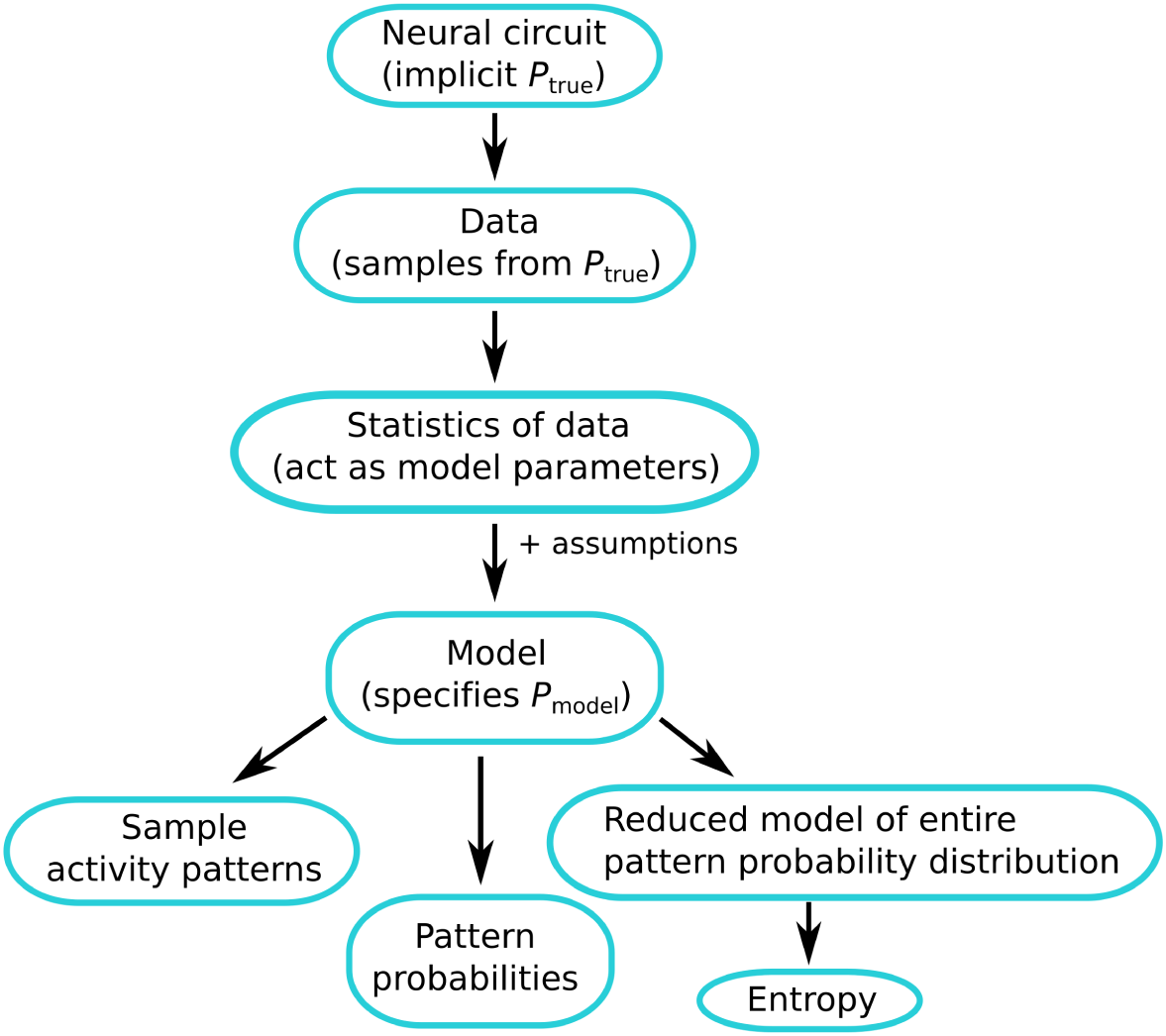
Schematic diagram of the model-building and utilization procedure. The neural circuit generates activity patterns sampled from some implicit distribution *P_true_*, which are recorded by an experimentalist as data. We estimate certain statistics of these data to be used as parameters for the model. The model is a mathematical equation that specifies a probability distribution over all possible patterns *P_moddel_*, whether or not each pattern was ever observed in the recorded data. We can then use the model for several applications: to sample synthetic activity patterns, to directly estimate pattern probabilities, or to build an even simpler model of the entire pattern probability distribution to estimate quantities such as the entropy.

The statistical model we propose for neural population data contains two sets of parameters that are fit in turn. The first set are the *N* free parameters needed to describe the population synchrony distribution: the probability distribution Pr(*K* = *k*) = *p*(*k*) for the number of neurons simultaneously active *K*, where 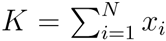. This distribution acts as a measure of the aggregate higher-order correlations in the population and so may contain information about the dynamical state of the network. For example, during network oscillations neurons may be mostly either all ON or all OFF together, whereas if the network is in an asynchronous mode, the population distribution will be narrowly centered around the mean neuron firing probabilities.

The second set of free model parameters are the conditional probabilities that each individual neuron is ON, given the total number of neurons active in the population, *p*(*x_i_* = 1|*K*). For shorthand we will write *p*(*x_i_*|*K*) instead of *p*(*x_i_* = 1|*K*) for the remainder of this paper. Since there are *N* + 1 possible values of *K*, and *N* neurons, there are *N*(*N* + 1) of these parameters. However, we know by definition that when *K* = 0 (all neurons are silent) and *K* = *N* (all neurons are active) then we must have *p*(*x*|*K* = 0) = 0 and *p*(*x*|*K* = *N*) = 1 respectively. Hence we are left with only *N*(*N* – 1) free parameters. Different neurons tend to have different dependencies on the population count, because of their heterogeneity in average firing rates (Buzsáki and Mizuseki, 2014) and because some neurons tend to be closely coupled to the activity of their surrounding population while others act independently (Okun et al., 2015). These two types of neurons have previously been termed ‘choristers’ and ‘soloists’, respectively.

Once the *N*^2^ total free parameters have been estimated from data (we discuss how this can be done below), we can construct the model. It gives the probability of seeing any possible activity pattern — even for patterns we have never observed — as

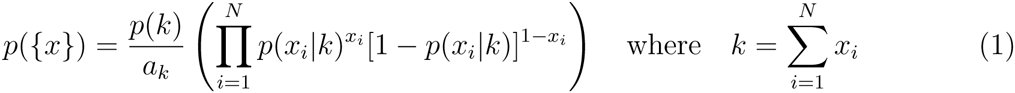

where *a_k_* is a normalizing constant defined as the sum of the probabilities of all 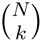 patterns in the set *S*(*k*) where 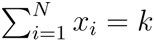 under a hypothetical model where neurons are conditionally independent:

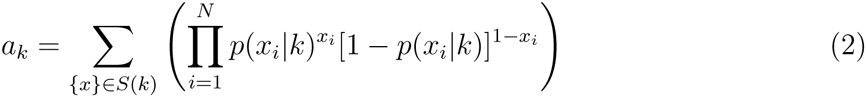

The model can be interpreted as follows: given the estimated synchrony distribution *p*(*k*) and set of conditional probabilities *p*(*x*_*i*_|*K*), we imagine a family of *N*–1 probability distributions *q_k_*({*x*}), *k* ∈ [1 : *N*–1] where pattern probabilities are specified by the conditional independence models 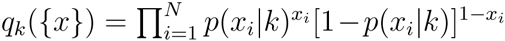. Now, using this family of distributions we construct one single distribution *p*({*x*}) by rejecting all patterns in each *q*_*k*_({_X_}) where 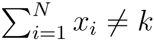, concatenating the remaining distributions (which cover mutually exclusive subsets of the pattern state space), and renormalizing so that the pattern probabilities sum to one. This implies that for any given activity pattern {*x*}, *p*({*x*}) ∝ *q_k_*({*x*}).

More intuitively, the model can be thought of as having two component ‘levels’: first, a high-level component that matches the distribution for the population rate. This component counts how many neurons are active, ignoring the neural identities and treating all neurons as homogeneous. The second, low-level component accounts for some of the heterogeneity between neurons. It asks, given a certain number of active neurons in the population, what is then the conditional probability that each individual neuron is active? This component captures two features of the data: the differences in firing rates between neurons, which can vary over many orders of magnitude (Buzsáki and Mizuseki, 2014), and the relationship between a neuron’s activity and the aggregate activity of its neighbors (Okun et al., 2015). Both of these features can potentially have large effects on the pattern probability distribution.

In Figure 2, we fit this statistical model to electrophysiology spike data recorded from a population of 50 neurons in macaque V1 while the animal was presented with a drifting oriented grating visual stimulus. A section of the original spiking data during stimulus presentation are shown in Figure 2A, top, along with synthetically generated samples from the model fitted to these data, below it in red. By definition the model matches the original data’s population synchrony distribution and conditional probability that each neuron is active (Figure 2B). In Figure 2C we show the model’s prediction for statistics of the data that it was not fitted to.

**Fig. 2:**
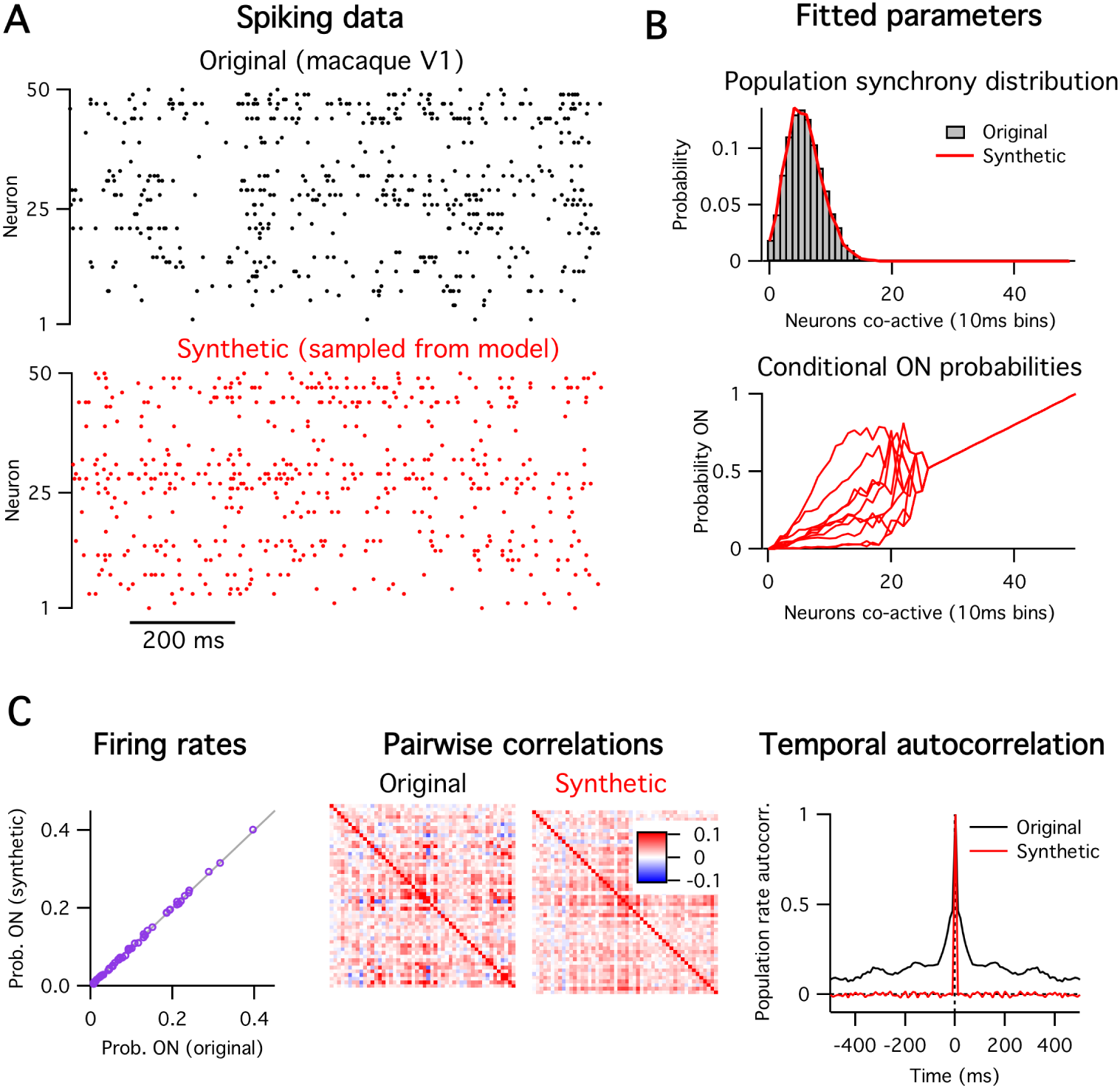
**A:** Original spiking data (top, black) and synthetic data generated from model (bottom, red). **B:** The model’s fitted parameters. First, the population synchrony distribution (top), and second the conditional probability that each neuron is ON given the number of neurons active. The conditional ON probabilities of only ten of the fifty neurons are shown for clarity. The curves converge to a straight line for *k* ≳ 25 because those values of *k* were not observed in the data, so the parameter estimates collapse to the prior mean. **C:** Comparison of other statistics of the data with the model’s predictions. The model gives an exact match of the single neuron firing rates (left), a partial match with the pairwise correlations (center), but does not match the data’s temporal correlations (right).

First (Figure 2C, left) the model almost exactly matches the average firing rate for each individual neuron. This is a direct consequence of the way the model is constructed and follows from the fits of the two sets of parameters. Hence the model can captures the heterogeneity in neural firing rates.

Second, we compare the pairwise correlations between neurons from the original data with those from the data synthetically generated by sampling the model (Figure 2C, center). Here we see only a partial match. Although the model captures the coarse features of the correlation matrix, it does not match the fine-scale structure on a pair-by-pair basis. For this example, the *R*^2^ value between the model and data pairwise correlations was 0.52 (Appendix Figure 1). In particular, the model accounts exactly for the population’s mean pairwise correlation, because this is entirely due to the fluctuations in the population activity. We can demonstrate this effect directly by first subtracting away the covariance in the original data that can be accounted for by the model and then renormalizing to get a new correlation matrix (Appendix Figure 1). Indeed this new correlation matrix is zero mean, but retains much of the fine-scale structure between certain pairs of neurons. This implies that the model captures only coarse properties of the pairwise correlations.

Finally, the model does not match at all the temporal correlations present in the original data (Figure 2C, right), since it assumes that each time bin is interchangeable. Note that this limitation is an ingredient of the model, not a failing *per se.* This property is shared with many other statistical methods commonly applied to neural population data (Schneidman et al., 2006; Macke et al., 2009; Cunningham and Yu, 2014; Okun et al., 2015).

These results show which statistics of the data that the population tracking model does and does not account for. Although other statistical models may more accurately account for pairwise or temporal correlation structure in the data, they typically do not scale well to large *N* (Table 1). In the remainder of the paper we explore the model’s behavior on large *N* data, and show how we can take advantage of the particular form of the model to robustly estimate some high-level measures of the activity statistics, including the entropy of the data and the divergence between pairs of data sets. Since these measures are typically difficult or impossible to estimate using other common statistical models in the field, the population tracking model may allow experimenters to ask neurobiological questions that would be otherwise intractable.

### 2.2 Fitting the model to data

We now outline a procedure for fitting the statistical model’s *N*^2^ free parameters to neural population data. We assume that the data have already been preprocessed as discussed above and are in the format of either a binary *N* × *T* matrix *M*, or as a two-column integer list of active timepoints and their associated neuron IDs. We found that parameter fitting was fast; for example, fitting parameters to data from a one hour recording of 140 neurons was done on a standard desktop in ∼ 1 minute.

#### 2.2.1 Fitting the population activity distribution

The first set of parameters are the *N* values specifying the probability distribution for the number of neurons active *p*(*k*). In principle *K* can take on any of the *N* + 1 values from 0 (the silent state) to *N* (the all ON state), but since we have the constraint that the probability distribution must normalize to one, one parameter can be calculated by default so we need only fit *N* free parameters to fully specify the distribution. The most straightforward way to do this is by histogramming, which gives the maximum likelihood parameter estimates. We simply count how many neurons are ON at each of the *T* timepoints to get [*K*(*t* = 1), *K*(*t* = 2)…*K*(*t* = T)], then histogram this list and normalize to one so that our estimate 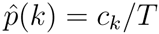 where *c_k_* is the count of the number of timepoints where k neurons were active.

If the data statistics are sufficiently stationary relative to the timescale of recording, then the error on each parameter individually scales 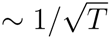 and independent of *N*. However, the relative error on each 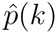 also scales 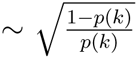, which implies large errors for rare values of *K*, when *p*(*k*) is small. Since neural activity is often sparse, we expect it to be quite common to observe small *p*(*k*) for large *K*, close to *N* (neurons are rarely all ON together). To avoid a case where we naively assign a probability of zero to a certain *p*(*k*) just because we never observe it in our finite data, we propose adding some form of regularization on the distribution *p*(*k*). A common method for regularization is to assume a prior distribution for *p*(*k*), then multiply it with the likelihood distribution from the data to compute the final posterior estimate for the parameters following Bayes rule. If for convenience we assume a Dirichlet prior (conjugate to the multinomial distribution), then the posterior mean estimate for each parameter simplifies to

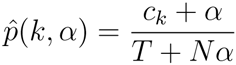

where *α* is a small positive constant. Note that this procedure is equivalent to adding the same small artificial count α to each empirical count *c_k_*. For the examples presented in this study, we set *α* = 0.01.

#### 2.2.2 Fitting the conditional ON probabilities for each neuron

The second step is to fit the *N*^2^ — *N* unconstrained conditional probabilities that each neuron is ON given the total number of active neurons in the population, *p*(*x|K*). The simplest method to fit these parameters is by histogramming, similar to the above case for fitting the population activity distribution. In this case we cycle through each value of *K* from 1 to *N*–1, find the subset of timepoints at which there were exactly *k* neurons active, and count how many times each individual neuron was active at those timepoints, *d_i, k_*. The maximum likelihood estimate for the conditional probability of the *i*th neuron being ON given k neurons in the population active is just 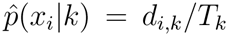, where *T_k_* is the total number of timepoints where *k* neurons were active.

As before, given that some values of *K* are likely to be only rarely observed we should also add some form of regularization to our estimates for *p*(*x|K*). We want to avoid erroneously assigning *p*(*x_i_*|*K*) = 0, or any *p*(*x_i_*|*K*) = 1 just because we had few data points available. Since *x_i_* here is a Bernoulli variable, we regularize following standard Bayesian practice by setting a Beta prior distribution over each *p*(*x_i_*|*K*) because it is conjugate to the binomial distribution. Under this model the posterior mean estimate for the parameters are

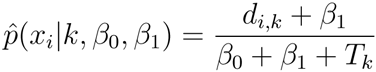

Using the Beta prior comes at the cost of setting its two hyperparameters, β_1_ and β_2_. We eliminate one of these free hyperparameters by constraining the prior’s mean to be equal to *k/N*. This will pull the final parameter estimates towards the values that they would take if all neurons were homogeneous. The other free hyperparameter is the variance or width of the prior. This dictates how much the final parameter estimate should reflect the data: the wider the prior is, the closer the posterior estimate will be to the naive empirical data estimate. We found in practice good results if the variance of this prior scaled with the variance of the Bernoulli variables, ∝ μ(1 − μ) where μ = *k/N*. This guaranteed that the variance vanished as *k* became near 0 or *N*. For the examples presented in this study, we set the prior variance *σ*^2^ = 0.5μ(1 — μ), and 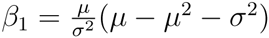, and 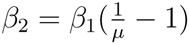.

An alternative method for fitting *p*(*x|K*) would be to perform logistic regression. Although in principle logistic regression should work well since we expect *p*(*x|K*) to typically be both monotonically increasing and correlated across neighboring values of *k*, we found in practice that as long as sufficient data were available it gave inferior fits compared with the histogram method discussed above. However for data sets with limited time samples logistic regression might indeed be preferable. The other benefit would be that since logistic regression requires fitting of only two parameters per regression, if employed it would reduce the total number of the model’s free parameters from *N*^2^ to only 3*N*.

#### 2.2.3 Calculating the normalization constants

The above expression for pattern probabilities includes a set of *N*–1 constants *A_k_* = {*a*_1_, *a*_2_ … *a*_*N*−1_} that are necessary to ensure that the distribution sums to one. These constants are not fit directly from data but instead follow from the parameters.

Each *a_k_* is calculated separately for each value of k. They can be calculated in at least four ways. The most intuitive method is via the brute force enumeration of the probabilities of all 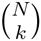 possible patterns where *k* neurons are active, then summing the probabilities, as given by eq. 2. Although this method is exact, it is only computationally feasible if 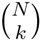 is not too large, which can occur quite quickly when analyzing data from more than 20–30 neurons. The second method to estimate *a_k_* is to draw *N* Bernoulli samples for many trials following the probabilities given by *p*(*x|k*), then count the fraction of trials in which the number of active neurons did in fact equal *k*. This method is approximate and inaccurate for large *N* because *a_k_* → 0 as *N* → ∞.

The third method is to estimate *a_k_* using importance sampling. We can rewrite eq. 2 as

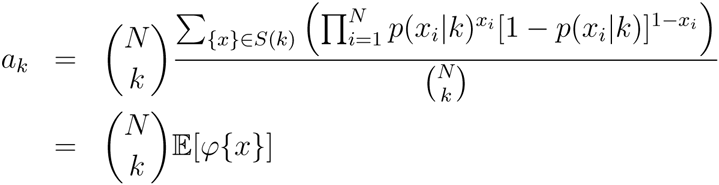

where {*x*} is a sample from the uniform distribution on *S*(*k*), and 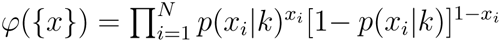. If we have m such samples {*x*^(1)^}, {*x*^(2)^}, …, {*x*^(m)^}, then by the law of large numbers

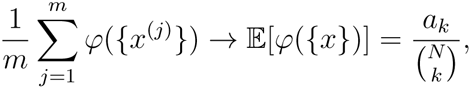

so by implication

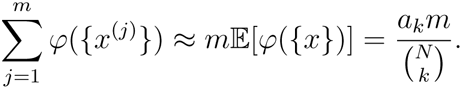

If we fit a straight line in *m* to the partial sums 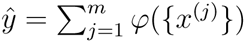 by linear regression, say 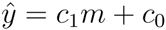, we get

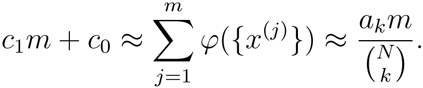

Assuming that 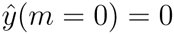, then the intercept c_0_ = 0, so we are left with

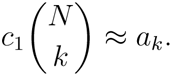

Finally, a fourth method follows from a procedure we present below, for estimating a low-dimensional model of the entire pattern probability distribution as a sum of log-normals.

#### 2.2.4 The implicit prior on the pattern probability distribution

By assuming a prior distribution over all of our parameters, we are implicitly assuming a prior distribution over the model’s predicted pattern probabilities. What does that look like? For the population activity distribution we have chosen a uniform value of *a* across all values of *k*, implying that our prior expects each level of population activity to be equally likely. The prior imposed on the second set of parameters, the *p*(*x|K*)’s, would assign each neuron an identical conditional ON probability of *k/N*. Although the second set of priors is maximal entropy given the first set, it is important to note that the uniform prior over population activity is not maximum entropy, since each value of *k* carries a different number of patterns. Hence for large N, the prior will be concentrated on patterns where few (*k* near zero) or many (*k* near *N*) neurons are active.

A geometrical view of the effect of the priors can be given as follows. Since our *N*^2^ parameters can each be written as a weighted linear sum of the 2^*N*^ pattern probabilities, they specify *N*^2^ constraint hyperplanes for the solution in the 2^*N*^-dimensional space of pattern probabilities. There are also other constraint hyperplanes which follow from constraints inherent to the problem, such as the fact that the pattern probabilities must sum to one, and that *p*(*x*|*K* = 0) = 0, etc. Since *N*^2^ < 2^*N*^ (for all *N* > 4) there are an infinite number of solutions that satisfy the constraints. Our final expression for the pattern probabilities is just a single point on the intersection of this set of hyperplanes. The effect of including priors on the parameters is to shift the hyperplanes so that our final solution is closer to prior pattern probabilities than that directly predicted by the data. In doing so it ensures all patterns are assigned a non-zero probability of occurring, as any sensible model should.

### 2.3 Sampling from model given parameters

Given the fitted parameters, sampling is straightforward using the following procedure:

1. Draw a sample for the integer number of neurons active *k_sample_* from the range {0,…, *N*} according to the discrete distribution *p*(*k*). This can be done by drawing a random number from the uniform distribution then mapping that value onto the inverse of the cumulative of *p*(*k*).
2. Draw *N* independent Bernoulli samples ***x*** = {*x*_1_, *x*_2_ … *x_N_*}, one for each neuron, with the probability for the *i*th neuron given by *p*(*x_i_*|*k_sample_*). This is a candidate sample.
3. Count how many neurons are active in the candidate sample: 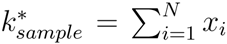. If 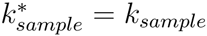, accept the sample. If 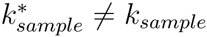, reject the sample and return to step 2.

One benefit of this model is that since the sampling procedure is not iterative, sequential samples are completely uncorrelated.

### 2.4 Estimating the full pattern probability distribution, entropy, and divergence

#### 2.4.1 Low-dimensional approximation to pattern probability distribution

So far we have shown how to fit the model’s parameters, calculate the probability of any specific population activity pattern, and sample from the model. Depending on the neurobiological question an experimenter might also wish to use this model to calculate the probabilities of all possible activity patterns, either to examine the shape of the distribution or to compute some measure that is a function of the entire distribution. One such measure, for example, is the joint population entropy H used in information theoretic calculations, 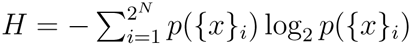.

For small populations of neurons *N* ≲ 20, the probabilities of all 2^*N*^ possible activity patterns can be exhaustively enumerated. However, for larger populations this brute force enumeration is not feasible due to limitations on computer storage space. For example, storing 2^100^ ∼ 10^30^ decimal numbers on a computer with 64-bit precision would require ∼ 10^19^ terabytes of storage space. Hence for most statistical models, such as classic pairwise maximum entropy models, this problem is either difficult or intractable (Broderick et al., 2007; although see Schaub and Schultz, 2012). Fortunately, the particular form of the model we propose implies that the distribution of pattern probabilities it predicts will, for sufficiently large *k* and *N*, tend towards the sum of a set of log-normal distributions, one for each value of *k* (Figure 3B–C), as we explain below. Since the log-normal distribution is specified by only 2 parameters, we can fit this approximate model with only 3N parameters total, which can be readily stored for any reasonable value of *N*.

**Fig. 3:**
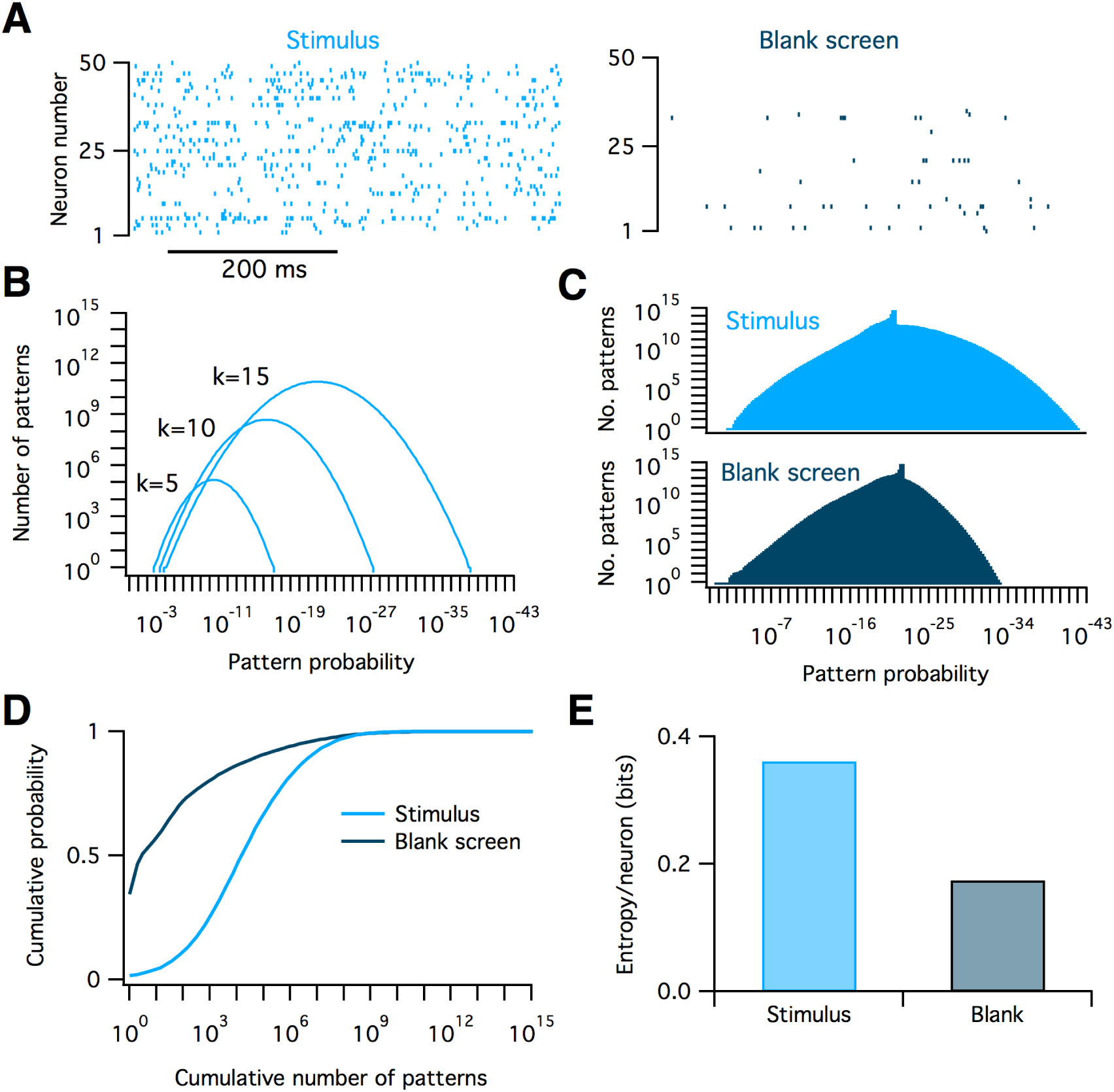
Calculating the distribution of population pattern probabilities and entropy for spiking data from macaque visual cortex. **A:** Example raster plots of spiking data from 50 neurons in macaque V1 in response to static oriented bar stimulus (left) and a blank screen (right). **B:** The distribution of pattern probabilities for varying numbers of neurons is estimated for various values of the numbers of neurons active, *k*. **C:** Summed total distribution of pattern probabilites for data recorded during stimulus (top, light blue) and blank screen (bottom, dark blue) conditions. The small bumps on top of the distributions are due to values of *k* which were unobserved in the data. Since the model assumes all patterns at these values are equally probable, they lead to the introduction of several sharp delta peaks to the pattern probability distribution. **D:** The cumulative probability as a function of the cumulative number of patterns considered. Note that many-fold fewer activity patterns account for the bulk of the probability mass in the blank screen condition compared to during the stimulus. **E:** Entropy per neuron of the pattern probability distribution for both conditions.

We derive the sum-of-lognormals distribution model as follows. First we take the log of both sides of eq.1 to get:

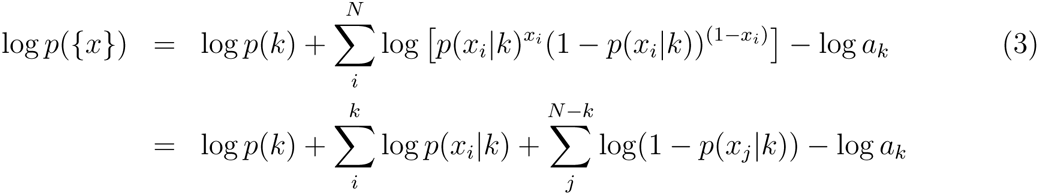

where the second and third terms correspond to sums over the *k* active and (*N* − *k*) inactive neurons in {*x*} respectively. Note that this equation is only valid for the cases where *k* ≥ 1. For clarity in what follows, we will temporarily represent *p*({*x*}) = θ and *p*({*x*}|*k*) = *θ_k_*. Now let us consider the set *L_k_* of the log-probabilities for all 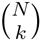 patterns for for a given level of population activity *k*, *L_k_* = {log(*p*({*x*})}_*k*_ = {log(θ)}_*k*_ where 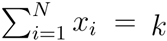. Since the population tracking model assumes that neurons are (pseudo) conditionally independent, then for sufficiently large N, according to the central limit theorem the second and third terms in the sum in eq. 3 will be normally distributed with some mean *μ*(*k*) and variance σ^2^(*k*), no matter what the actual distribution of *p*(*x*_*i*_|*K*)’s is. Hence, if we were to histogram the log-probabilities {log(θ)}_*k*_ of all patterns for a given k, their distribution could be approximated by the sum of two Gaussians and two constants:

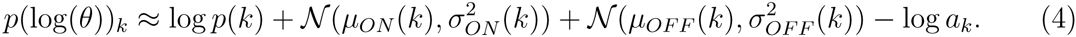

Note that this is a distribution over log–pattern probabilities: it specifies the fraction of all neural population activity patterns that share a particular log–probability of being observed.

The two normal distribution means are given by

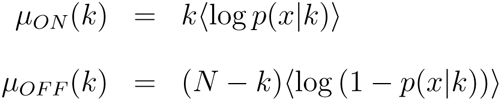

and the variances are

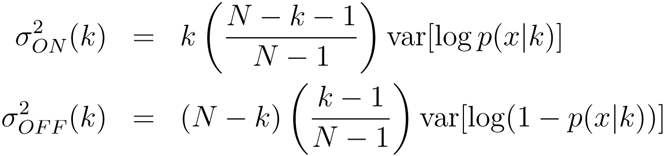

where the fractional terms in the variance equations are corrections because we are drawing without replacement from a finite population. Finally since we are adding two random variables (the second and third terms in 4), we also need to account for their covariance. Unfortunately, the value of this covariance depends on the data, and unlike the means and variances we could find no simple formula to predict it directly from the parameters *p*(*x|k*). Hence it should be estimated empirically by drawing random samples from the coupled distributions 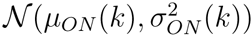 and 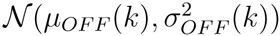, and computing the covariance of the samples.

Although the lognormal approximation is valid when both *K* and *N* are large, the approximation will become worse when *K* is near 0 and N, no matter how large *N* is. This is problematic because neural data is often sparse, so small values of *K* are expected to be common and hence important to accurately model. Indeed we found empirically that the distribution of log-pattern probabilities at small *K* can become substantially skewed, or, if the data come from neurons that include distinct subpopulations with different firing rates, even multimodal. We suggest that the experimenter examines the shape of the distribution by histogramming the probabilities of a large number of randomly chosen patterns to assess the appropriateness of the lognormal fit. The validity of the log-normal approximation can be formally assessed using, for example, the Lilliefors or Anderson-Darling tests. If the distribution is indeed non-lognormal for certain values of *K*, we suggest application of either or both of the following two *ad hoc* alternatives. First, for very small values of *K* (say *k* ≲ 3), then the number of patterns at this level of population synchrony 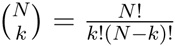 should also be small enough to permit brute force enumeration of all such pattern probabilities. Second, for slightly larger values of *K* (3 ≲ *k* ≲ 10), the distribution can be empirically fit by alternative low-dimensional parametric models, for example a mixture-of-gaussians (MoG), which should be sufficiently flexible to capture any multimodality or skewness. In practice we found that MoG model fits are typically improved by initializing the parameters with standard clustering algorithms, such as K-means.

One important precaution to take when fitting any parametric model to the pattern probability distributions (be it lognormal, MoG, or otherwise) is to make sure that the resulting distributions are properly normalized so that the product of the integral of the approximated distribution of pattern probabilities for a given *k*, *p*(θ)_*k*_, with the total number of possible patterns at that *k*, 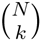, does indeed equal the *p*(*k*) previously estimated from data:

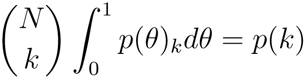

Although in principle this normalization should be automatic as part of the fitting procedure, even small errors in the distribution fit due to finite sampling can lead to appreciable errors in the normalization, due to the exponential sensitivity of the pattern probability sum on the fit in log co-ordinates. The natural place to absorb this correction is in the constant *a_k_*, which in any case has to be estimated empirically so it will carry some error. Hence we suggest that when performing this procedure, estimation of *a*_k_ should be left as the final step, when it can be calculated computationally as whatever value is necessary to satisfy the above normalization.

#### 2.4.2 Calculating population entropy

Given the above reduced model of the pattern probability distribution we could compute any desired function of the pattern probabilities, for example the mean or median pattern probability, the standard deviation, etc. One example measure that is relevant for information theory calculations is Shannon’s entropy, H = − Σ_*i*_*p_i_* log2*p_i_*, measured in bits. This can be calculated by first decomposing the total entropy as

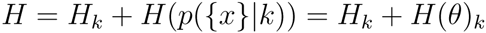

where 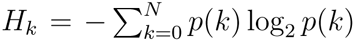 is the entropy of the population synchrony distribution and 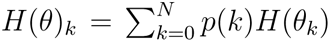 is the conditional entropy of the pattern probability distribution given *K*. Given the sum-of-lognormals reduced model of the pattern probability distribution, the total entropy (in bits) of all patterns for a given *k* is

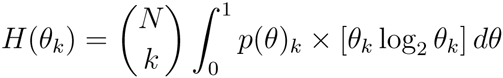

This can be calculated by standard numerical integration methods separately for each possible value of *K*.

In the homogeneous case where all neurons are identical, all 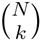 patterns for a given *K* will have equal probability of occurring, 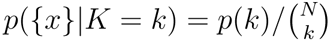. This situation maximizes the second term in the entropy expression, and simplifies it to 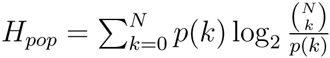.

To demonstrate these methods we calculated the probability distribution across all 2^50^ ≈ 10^15^ possible population activity patterns, and the population entropy, for an example spiking data set recorded from fifty neurons in macaque primary visual cortex. The presentation of a visual stimulus increases the firing rates of most neurons as compared to a blank screen (Figure 3A). We found that this increase in firing rates lead to a shift in the distribution of pattern probabilities (Figure 3C–D) and an increase in population entropy (Figure 3E). Notably, a tiny fraction of all possible patterns account for almost all the probability mass. For the visually evoked data, around 10^7^ patterns accounted for 90% of the total probability, which implies that only 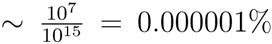 of all possible patterns are routinely used. Although this result might not seem surprising given that neurons fire sparsely, any model that assumed independent neurons would likely overestimate this fraction because such a model would also overestimate the neural population’s entropy (see below). These results demonstrate that the population tracking model can detect aspects of neural population firing that may be difficult to uncover with other methods.

#### 2.4.3 Calculating the divergence between model fits to two data sets

Many experiments in neuroscience involve comparisons between neural responses under different conditions: for example the firing rates of a neural population before and after application of a drug, or the response to a sensory stimulus in the presence or absence of optogenetic stimulation. Therefore it would be desirable to have a method for quantifying the differences in neural population pattern probabilities between two conditions. Commonly used measures for differences of this type are the Kullback-Leibler divergence, and the related Jensen-Shannon divergence (Cover and Thomas, 2006; Berkes et al., 2011). Calculation of either divergence involves a point-by-point comparison of the probabilities of each specific pattern under the two conditions. For small populations, this can be done by enumerating the probabilities of all possible patterns, but how would it work for large populations? On the face of it, the above approximate method for entropy calculation cannot help here, because that involved summarizing the distribution of pattern probabilities while losing the identities of individual patterns along the way. Fortunately the form of the statistical model we propose does allow for an approximate calculation of the divergence between two pattern probability distributions, as follows.

The Kullback-Leibler divergence from one probability distribution *p*(*i*) to another probability distribution *q*(*i*) is defined as

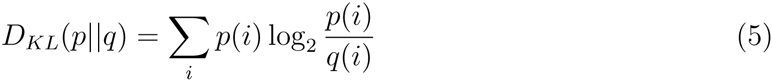

We can decompose this sum into *N* + 1 separate sums over the subsets of patterns with *K* neurons active:

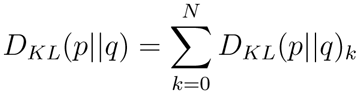

Hence we just need a method to compute *D_KL_*(*p*‖*q*)_*k*_ for any particular value of *k*. Notably, the term to be summed over in equation 5 can be seen as the product of two components: *p*(*i*) and 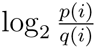. In the preceding section we showed that for sufficiently large *k* and N, the distribution of pattern probabilities at a fixed *K* is approximately log-normal because of our assumption of conditional independence between neurons. Hence the first component *p*(*i*) can be thought of as a continuous random variable that we will denote *X_1_*, drawn from the log-normal distribution *f*(*x*_1_). Because *p*(*i*) represents pattern probabilities, the range of *f*(*x*_1_) is [0,1]. The second component, 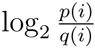, in contrast, can be thought of as a continuous random variable that we will denote *X*_2_, that is drawn from the normal distribution *g*(*x*_2_), because by the same argument 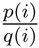 is approximately log-normally distributed, so its logarithm is normally distributed. Since this term is the logarithm of the ratio of two positive numbers, the range of *g*(*x*_2_) is [—∞, ∞]. Now the term to be summed over can be thought of as the product of two continuous and dependent random variables *Y* = *X*_1_*X*_2_, with some distribution *h*(*y*).

Our estimate for the KL divergence 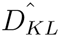 for a given *k* is then just the number of patterns at that value of *k* times the expected value of *Y*:

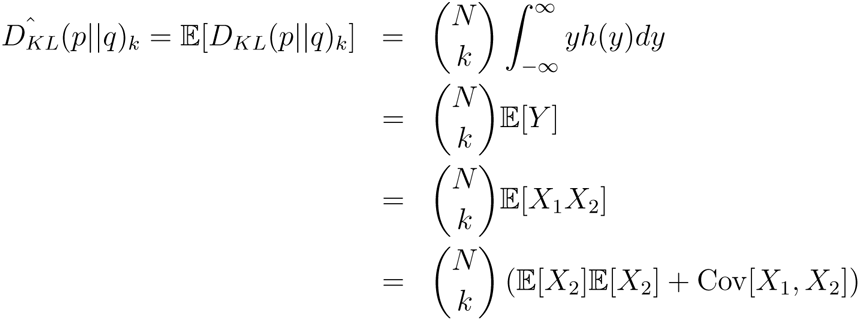

The three new terms in the last expression, 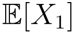, 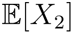, and Cov[*X*_1_, *X*_2_], can be estimated empirically by sampling a set of matched values of *p*({*x*}_*i*_) and *q*({*x*}_*i*_) from a large randomly chosen subset of the 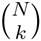 patterns corresponding to a given value of *k*.

### 2.5 Model fit convergence for large numbers of neurons

To test how the model scales with numbers of neurons and time samples, we fit it to synthetic neural population data from a different established statistical model, the Dichotomized Gaussian (DG) (Macke et al., 2009). The DG model generates samples by thresholding a multivariate Gaussian random variable in such a way that the resulting binary values matches desired target mean ON probabilities and pairwise correlations. The DG is a particularly suitable model for neural data, because has been shown that the higher-order correlations between ‘neurons’ in this model reproduce many of the properties of high-order correlations seen in real neural populations recorded *in vivo* (Macke et al., 2011b). This match may come from the fact that thresholding behavior of the DG model mimics the spike threshold operation of real neurons.

For this section we used the DG to simulate the activity of two equally sized populations of neurons, *N*_1_ = *N*_2_ = *N*/2, one population with a low firing rate of *r*_1_ = 0.05 and the other with a higher firing rate of *r*_2_ = 0.15. The correlations between all pairs of neurons were set at *ρ* = 0.1. We first estimated ground truth pattern probability distributions by histogramming samples. Although there are 2^*N*^ possible patterns, the built-in symmetries in our chosen parameters meant that all patterns with the same number of neurons active from each group *k*_1_ and *k*_2_ share identical probabilities. Hence the task amounted to estimating only the joint probabilities *p*(*k*_1_, *k*_2_) of the (*N*+1)^2^ configurations of having *k*_1_ and *k*_2_ neurons active. We generated as many time samples as was needed for this probability distribution to converge (T > 10^9^) for varying numbers of neurons ranging from *N* = 10 to *N* = 1000.

We then fit both our proposed model and several alternatives to further sets of samples from the DG, varying *T* from 100 to 1,000,000. Finally, we repeated the fitting procedure on many sets of fresh samples from the DG to examine variability in model fits across trials. To assess the quality of the fits we use the population entropy as a summary statistic. We compared the entropy estimates of the population tracking model with five alternatives:

1. Independent neuron model: neurons are independent, with individually fit mean firing rates estimated from the data. This model has *N* parameters.
2. Homogeneous population model: neurons are identical but not independent. The model is constrained only by the population synchrony distribution *p*(*k*), as estimated from data. This model has *N* + 1 parameters.
3. Histogram. The probability of each population pattern is estimated by the counting the number of times it appears and normalizing by *T*. This model has 2^*N*^ parameters.
4. Singleton entropy estimator (Berry II et al., 2013): this model uses the histogram method to estimate the probabilities of observed patterns in combination with an independent neuron model for the unobserved patterns. We implemented this method using our own MATLAB code.
5. Archer-Park-Pillow (APP) method (Archer et al., 2013): a Bayesian entropy estimator that combines the histogram method for observed patterns with a Dirichlet prior constrained by the population synchrony distribution. We implemented this method using the authors’ publicly available MATLAB code (http://github.com/pillowlab/CDMentropy).

We chose these models for comparison because they are tractable to implement. Although it is possible that other statistical approaches such as the maximum entropy model family would more accurately approximate the true data distribution, it is difficult to estimate the joint entropy from these models for data from ≳ 20 neurons (Table 1).

In Figure 4 we plot the mean and standard deviation of the entropy/neuron estimates for this set of models as a function of the number of neurons (panels B and C) and number of time samples (panels D and E) analyzed. The key observation is that across most values of *N* and *T*, the majority of methods predict entropy values different from the true value (dashed line in all plots). These errors in the entropy estimates come from three sources: the finite sample variance, the finite sample bias and the asymptotic bias.

**Fig. 4:**
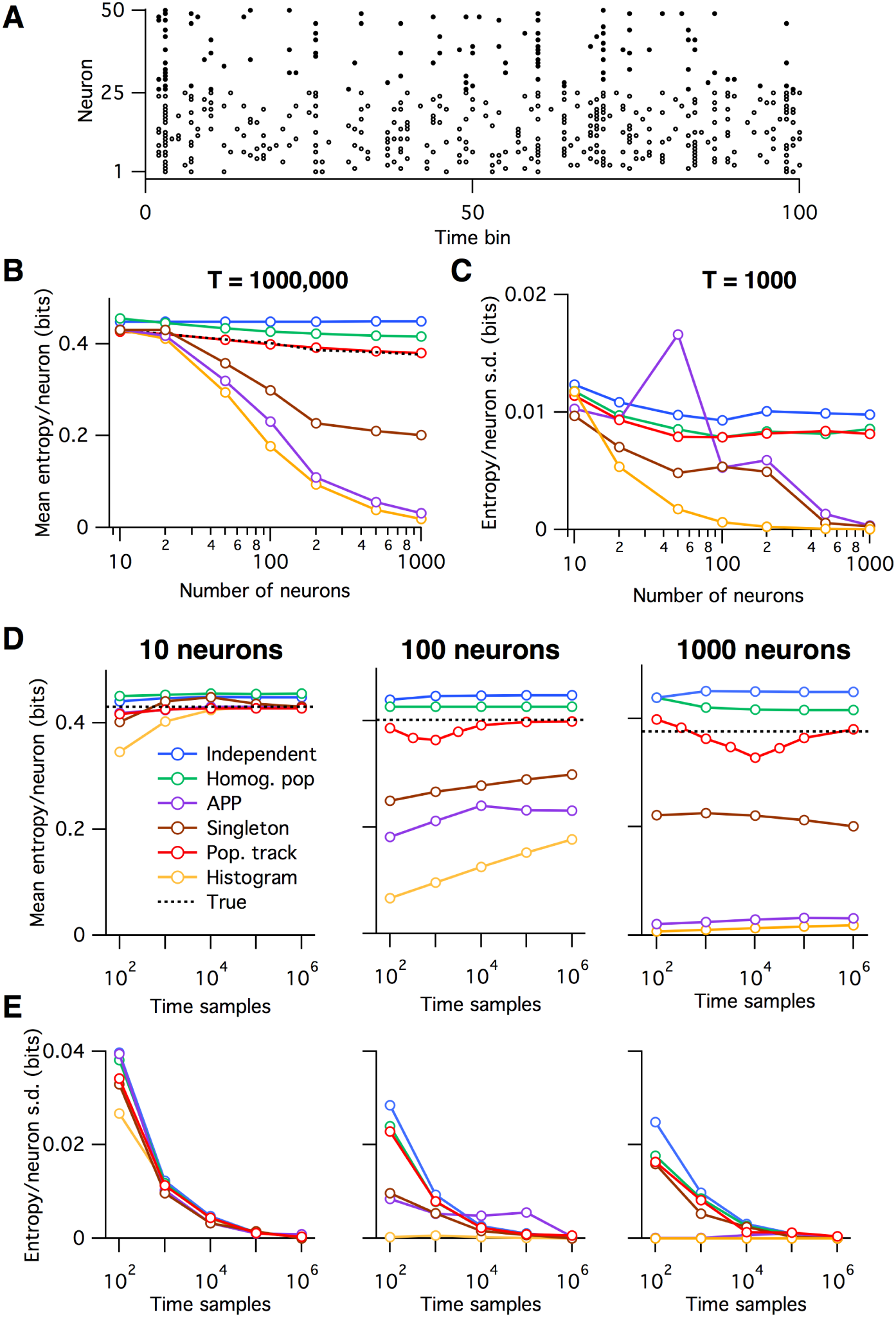
Convergence of entropy estimate as a function of the number of neurons and time samples analyzed. **A:** Example spiking data from the DG model with two subpopulations, a low firing rate group (filled black circles) and a higher firing rate group (open circles). **B-C:** Mean (B) and standard deviation (C) of estimated entropy per neuron as a function of the number of neurons analyzed, for each of the various models. **D-E:** The mean (D) and standard deviation (E) of estimated entropy per neuron as a function of the number of timesteps considered, for data from varying numbers of neurons (left to right).

The finite sample variance is the variability in parameter estimates across trials from limited data, shown in Figure 4C and E as the standard deviation in entropy estimates. Notably, the finite sample variance decreases to near zero for all models within 10^5^–10^6^ time samples, and is approximately independent of the number of neurons analyzed for the population tracking method (Figure 4C and E).

The second error, the finite sample bias, arises from the fact that entropy is concave function of *p*({*x*}). This bias is downward in the sense that the mean entropy estimate across finite-data trials will always be less than the true entropy: 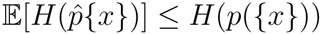. Intuitively, any noise in the parameter estimates will tend to make the predicted pattern probability distribution more lumpy than the true distribution, so reducing the entropy estimate. Although this error becomes negligible for all models within a reasonable number of time samples for small numbers of neurons (N ≈ 10) (Figure 4B and D), it introduces large errors for the histogram, singleton and APP methods for larger populations. In contrast to the finite sample variance, the finite sample bias depends strongly on the number of neurons analyzed for all models, typically becoming worse for larger populations.

The third error, the asymptotic bias, is the error in entropy estimate that would persist even if infinite time samples were available. It is due to a mismatch between the form of the statistical model used to describe the data and the true underlying structures in the data. In Figure 4, this error is present for all models that do not include a histogram component: the independent, homogeneous population and population tracking models. Because the independent and homogeneous population models are maximum entropy given their parameters, their asymptotic bias in entropy will always be ‘upward’, meaning that these models will always overestimate the true entropy, given enough data. They are too simple to capture all of the structure in the data. Although population tracking method may have either an upward or downward asymptotic bias, depending on the structure of the true pattern probability distribution, for the example cases we examined this error was small in magnitude.

The independent, homogeneous population, and population tracking models converged to their asymptotic values within 10^4^–10^5^ time samples (Figure 4D–E). The histogram, singleton and APP methods, in contrast, performed well for small populations of neurons, *N*< 20, but strongly underestimated the entropy for larger populations (Figure 4B, D), even for *T* = 10^6^ samples.

The independent, homogeneous population and population tracking models consistently predicted different values for the entropy. In order from greatest entropy to least entropy, they were: independent model, homogeneous population model, and population tracking. Elements of this ordering are expected from the form of the models. The independent model matches the firing rate of each neurons but assumes that they are uncorrelated, implying a high entropy estimate. Next, we found that the homogeneous population model had lower entropy than the independent model. However this ordering will depend on the statistics of the data so may vary from experiment to experiment. The model we propose, the population tracking model, matches the data statistics of both the independent model and the homogeneous population model. Hence its predicted entropy must be less than or equal to both of these two previous models. One important note is that the relative accuracies of the various models should not be taken as fixed, but will depend both on the statistics of the data and on the choices of the priors.

In summary, of the six models we tested on synthetic data, the population tracking model consistently performed best. It converged on entropy estimates close to the true value even for data from populations as large as 1000 neurons.

### 2.6 Population tracking model accurately predicts probabilities for both seen and unseen patterns

The above analysis involved estimating a single summary statistic, the entropy, for the entire 2^*N*^-dimensional pattern probability distribution. But how well do the models do at predicting the probability of individual population activity patterns? To test this we fit four of the six models to the same DG-generated data as the previous section, with *N* = 100 and *T* = 10^6^. As seen in Figure 4D–E, for data of this size the entropy predictions of the three statistical models had converged, but the histogram method’s estimate had not. We then drew 100 new samples from the same DG model, calculated all four models’ predictions of pattern probability for each sample, and compared the predictions with the known true probabilities (Figure 5).

**Fig. 5:**
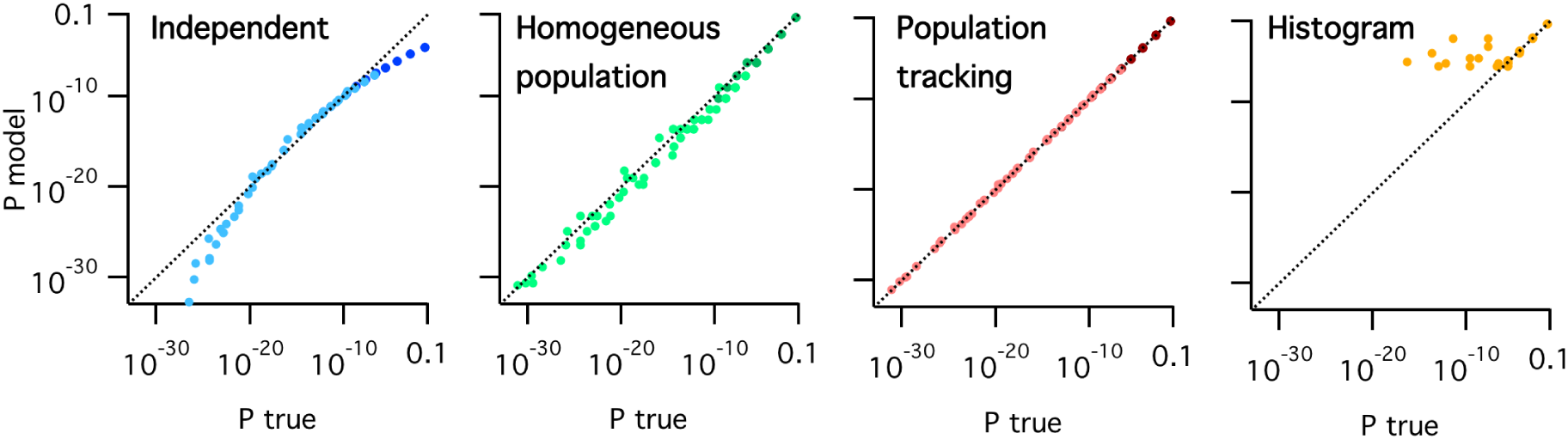
Predicted pattern probabilities as a function of true pattern probabilities for a population of 100 neurons sampled from the same DG model as Figure 4. From left to right: independent model (blue), homogeneous population model (green), population tracking model (red) and histogram method (amber). In each plot the darker colored symbols correspond to patterns seen during model training and so were used in fitting the model parameters, and lighter colored symbols correspond to new patterns that appeared only in the test set. The histogram plot (right) shows only data for the subset of patterns seen in both the training and test sets. Dashed diagonal line in each plot indicates identity.

The independent model’s predictions deviated systematically from the true pattern probabilities. In particular, it tended to underestimate both high-probability and low-probability patterns, while overestimating intermediate probability patterns. It is important to note that the data in Figure 5 are presented on a log scale. Hence these deviations correspond to many orders of magnitude error in pattern probability estimates. The homogeneous population model did not show any systematic biases in probability estimates but did show substantial scatter around the identity line, again implying large errors. This is to be expected since this model assumes that all patterns for a given *k* have equal probability. In contrast to these two models, the population tracking model that we propose accurately estimated pattern probabilities across the entire observed range. Finally, the histogram method failed dramatically. Although it predicted well the probabilities for the most likely patterns, it quickly deviated from the true values for more rare patterns. And worst of all, it predicts a probability of zero for patterns that it has not seen before, as evidenced by the large number of missing points in the right plot in Figure 5.

One final important point is that of the 100 test samples drawn from the DG model, 63 were not part of the training set (light colored circles in Figure 5). However, the population tracking model showed no difference in accuracy for these unobserved patterns compared with the 37 patterns previously seen during training (dark circles in Figure 5). Together, these results show that the population tracking model can accurately estimate probabilities of both seen and unseen patterns, for data from large numbers of neurons.

### 2.7 Model performance for populations with heterogeneous firing rates and correlations

In order to calculate the ground truth pattern probabilities and entropy for large *N* for the above analysis, we assumed homogeneous firing rates and correlations to ensure symmetries in the pattern probability distributions. However, since the population tracking model also implicitly assumes some shared correlations across neurons due to their shared dependence on the population rate variable *K*, this situation may also bias the results in favor of the population tracking model in the sense that this may be the regime where *P_model_* best matches *P_true_*. Since *in vivo* neural correlations typically appear to have significant structure (Figure 1C), we also examined the behavior of the model for a scenario with more heterogeneous firing rates and correlations. We repeated the above analysis using samples from the DG neuron model with *N* = 10, but with individual neuron firing rates drawn from a normal distribution *μ* = 0.1, *σ* = 0.02, and pairwise correlations drawn from a normal distribution with *μ* = 0.05, *σ* = 0.03 (Figure 6A). We numerically calculated the 2^10^ = 1024 ground truth pattern probabilities by exhaustively sampling from the DG model. We again varied the number of time samples from 100 to 1,000,000 and fit the population tracking model and several comparison models: the independent neuron model, the homogeneous population model, the histogram method, and also the pairwise maximum entropy model (Schneidman et al., 2006). We computed the Jensen-Shannon (JS) divergence, which is a measure of the difference between the true and model pattern probability distributions (Figure 6B), entropy/neuron (Figure 6C), and all 1024 individual pattern probabilities (Figure 6D). Although the population tracking model outperformed the independent and homogeneous population models as before, it was outperformed by the pairwise maximum entropy model on this task. The JS divergence of the population tracking model saturated at a higher non-zero floor than the pairwise maximum entropy models in Figure 6B. However, on the other hand the asymptotic error in the population tracking’s estimate of entropy was minimal at +0.0015 bits, or 0.3% (Figure 6C). It is difficult to ascertain whether the pairwise maximum entropy model would also outperform the population tracking model for large N, and requires further study.

**Fig. 6:**
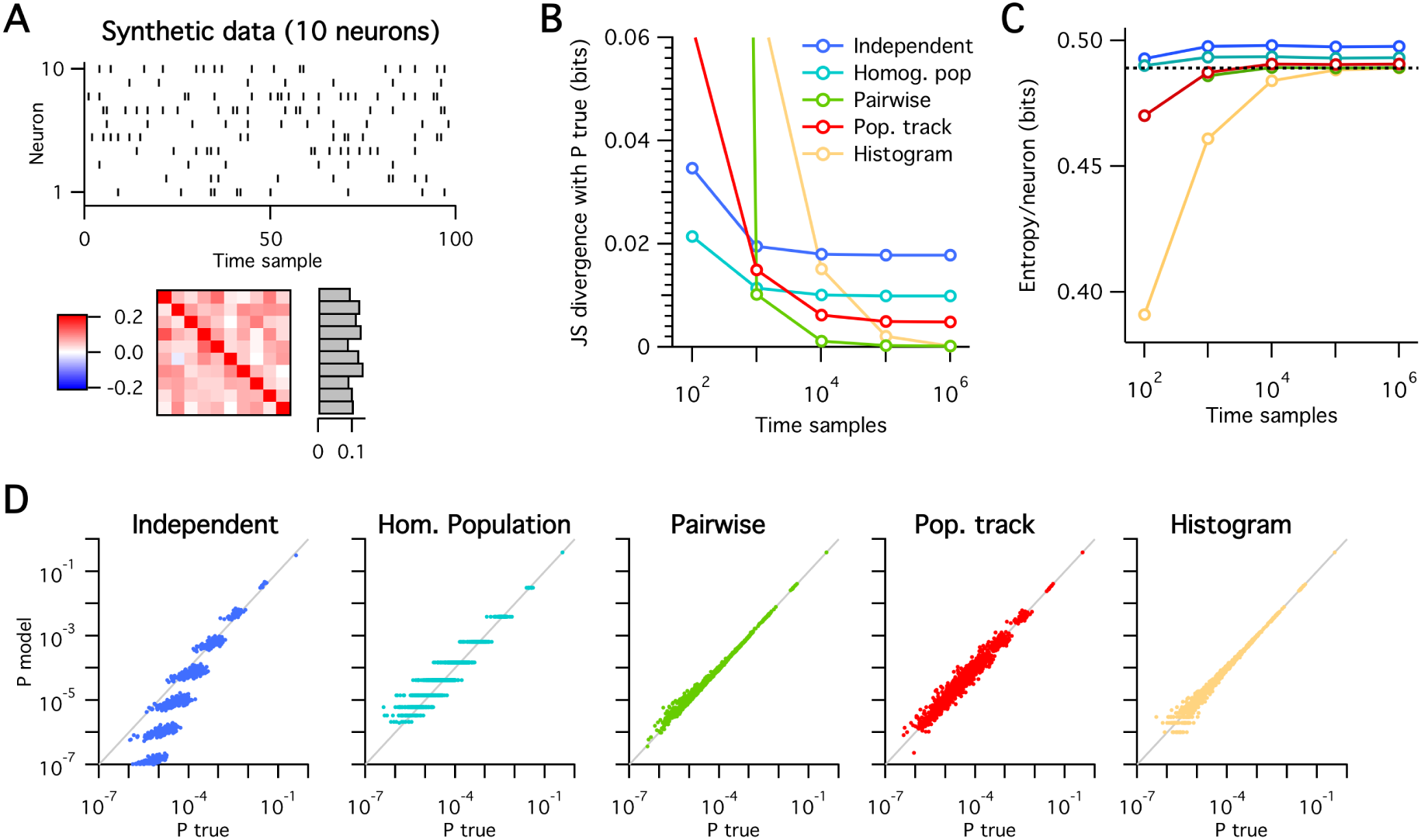
Performance of various models for data from 10 neurons with heterogeneous firing rates and correlations. **A:** Example spiking data from the DG model (top left), with heterogeneous correlations and firing rates (bottom). **B–C:** Jensen-Shannon divergence of each model’s predicted pattern probability distribution with the true distribution (B) and entropy per neuron (C) as a function of the number of time samples. **D:** Predicted pattern probabilities versus true pattern probabilities for each of the tested models (left to right), for 1,000,000 time samples.

### 2.8 Decoding neural population electrophysiological data from monkey visual cortex

We next tested the ability of the population tracking model to decode neural population responses to stimuli. We analyzed electrode array data recorded from anesthetized macaque primary visual cortex in response to visual stimuli (Figure 7A, see Experimental Procedures and Zandvakili and Kohn, 2015 for details). Spike sorting algorithms were applied to the raw voltage waveforms to extract the times of action potentials from multi-units. Altogether 131 different multi-units were recorded from a single animal. The animal was shown drifting oriented sinusoidal gratings chosen from eight orientations in a pseudorandom order. Each 1.28 s stimulus presentation was interleaved with a 1.5 s blank screen, and all eight possible stimulus orientations were presented 300 times each.

**Fig. 7:**
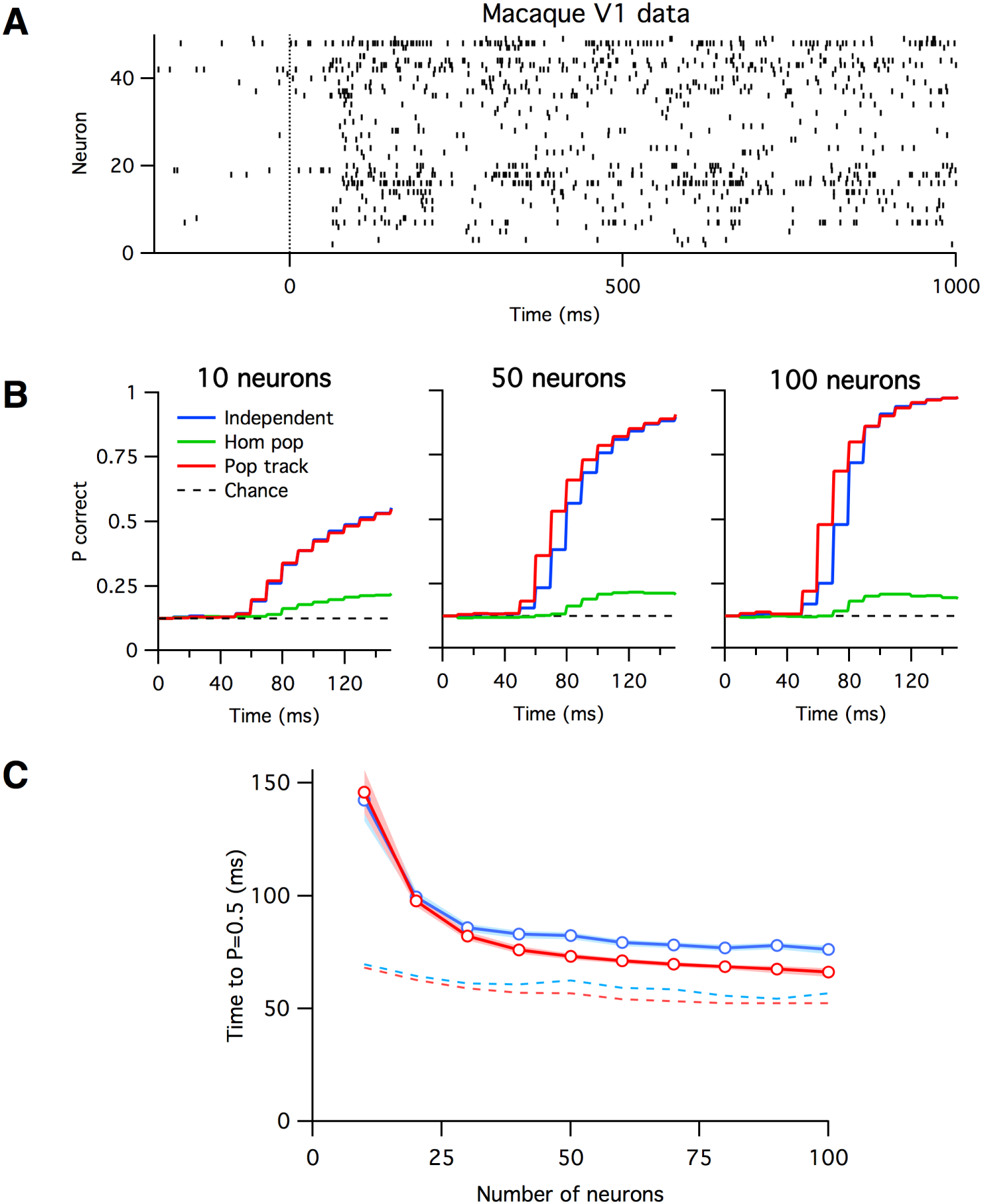
Decoding neural population spiking data from macaque primary visual cortex in response to oriented bar visual stimuli. **A:** Example spiking data from fifty neurons during a single presentation of an oriented bar stimulus. Time zero indicates onset of stimulus. **B:** Decoding performance as a function of time since stimulus onset for three different decoding models (different colored curves) and varying numbers of neurons (plots from left to right). Chance decoding level in all cases was 1/8 = 0.125. **C:** The mean time since stimulus onset to reach 50% decoding accuracy for the independent (blue) and population tracking (red) models, as a function of the number of neurons analyzed. The dashed curves indicate the time at which decoding accuracy first statistically exceeded noise levels. Time bin size fixed at 10 ms. The homogeneous population model is not shown because it never reached 50% decoding accuracy.

Our decoding analysis proceeded as follows. We first rebinned the data into 10 ms intervals. If a unit spiked one or more times in a time bin, it was labeled as ON, otherwise it was labelled OFF. Second, we chose a random subset of *N* units from the 131 total, and excluded data from the rest. Then for a given stimulus orientation, we randomly split the data from the 300 trials into a 200 trial training set, and 100 trial test set. We concatenated the data from the 200 training trials and fit the population tracking model to this dataset, along with two control statistical models: the independent model and the homogeneous population model. We repeated this procedure separately for the eight different stimulus orientations, so were left with eight different sets of fitted parameters, one for each orientation. We then applied maximum likelihood decoding separately on neural responses to 100 randomly chosen stimuli from the test dataset. Finally, we repeated the entire analysis 100 times for different random subsets of *N* neurons and training/test data set partitions, and took a grand average of decoding performance.

We plot the decoding performance of the various statistical models as a function of time since the stimulus onset in Figure 7B. For all models, decoding was initially at chance level (1/8 = 0.125), then began to increase around 50 ms after stimulus onset, corresponding to the delay in spiking response in visual cortex (Figure 7A). Decoding performance generally improved monotonically both with the number of neurons and number of timepoints analyzed, for all models. However, decoding performance was much higher for the independent and population tracking models, which saturated at almost 100% correct, compared with ∼ 25% correct for the homogeneous population model. Hence for these data it appears that the majority of information about the stimulus is encoded in the identities of which neurons are active, and not in the total numbers of neurons active.

Although both the independent and population tracking models saturated to almost 100% decoding performance at long times, we found that for larger sets of neurons, the population tracking model’s performance rose earlier in time than the independent neuron model (Figure 7B-C). For 10 neurons, the independent model and population tracking model reached 50% accuracy at similar times after stimulus onset (146 ms with 95% c.i. [136.4 : 156] ms for population tracking model and 142.5 ms with 95% c.i. [133.2 : 152.3] ms for independent model). However given spiking data from 100 neurons, the population tracking model reached 50% correct decoding performance at 66.1 ms after stimulus onset (95% c.i. [64.2 : 68] ms), whereas the independent model reached the same level later, at 76.2 ms after stimulus onset (95% c.i. [74.2 : 78] ms). Although superficially this may appear to be a modest difference in decoding speed, it is important to note that the baseline time for decoding above chance was not until 52.3 and 56.8 ms after stimulus onset for the population tracking and independent models, respectively (see Experimental Procedures for details). The reason for this late rise in decoding accuracy is the documented ∼ 50 ms lag in spiking response in macaque V1 relative to stimulus onset (Chen et al., 2006, 2008) (see Figure 7A). Given that we discretized the data into timebins of 10 ms, this implies that the population tracking model could decode stimuli mostly correctly given data from less 2 time frames on average. In summary, these results show that the population tracking model can perform rapid stimulus decoding.

### 2.9 Entropy estimation from two-photon Ca^2+^ imaging population data from mouse somatosensory cortex

As a second test case neurobiological problem, we set out to quantify the typical number of activity patterns and entropy of populations of neurons in mouse neocortex, across development. We applied our analysis method to spontaneous activity in neural populations from data previously recorded (Gonçalves et al., 2013) by in vivo two-photon Ca^2+^ imaging in layer 2/3 primary somatosensory cortex of unanesthetized wild-type mice with the fluorescent indicator Oregon green BAPTA-1 (see Experimental Procedures for further details). The original data were recorded at ∼ 4 Hz (256 ms timeframes), but for this analysis we resampled the data into 1 s timebins because we found that it optimized a tradeoff between catching more neurons in the active state versus maintaining a sufficient number of timeframes for robust analysis.

To compare neural activity across development we used the Shannon entropy/neuron, *h* (Figure 8H-I). Shannon entropy is a concept adopted from information theory that quantifies the uniformity of a probability distribution. If all patterns were equally probable then *h* = 1 bit. At the opposite extreme, if only one pattern were possible then *h* = 0 bits. It also has a functional interpretation as the upper limit on the amount of information the circuit can code (Cover and Thomas, 2006).

**Fig. 8:**
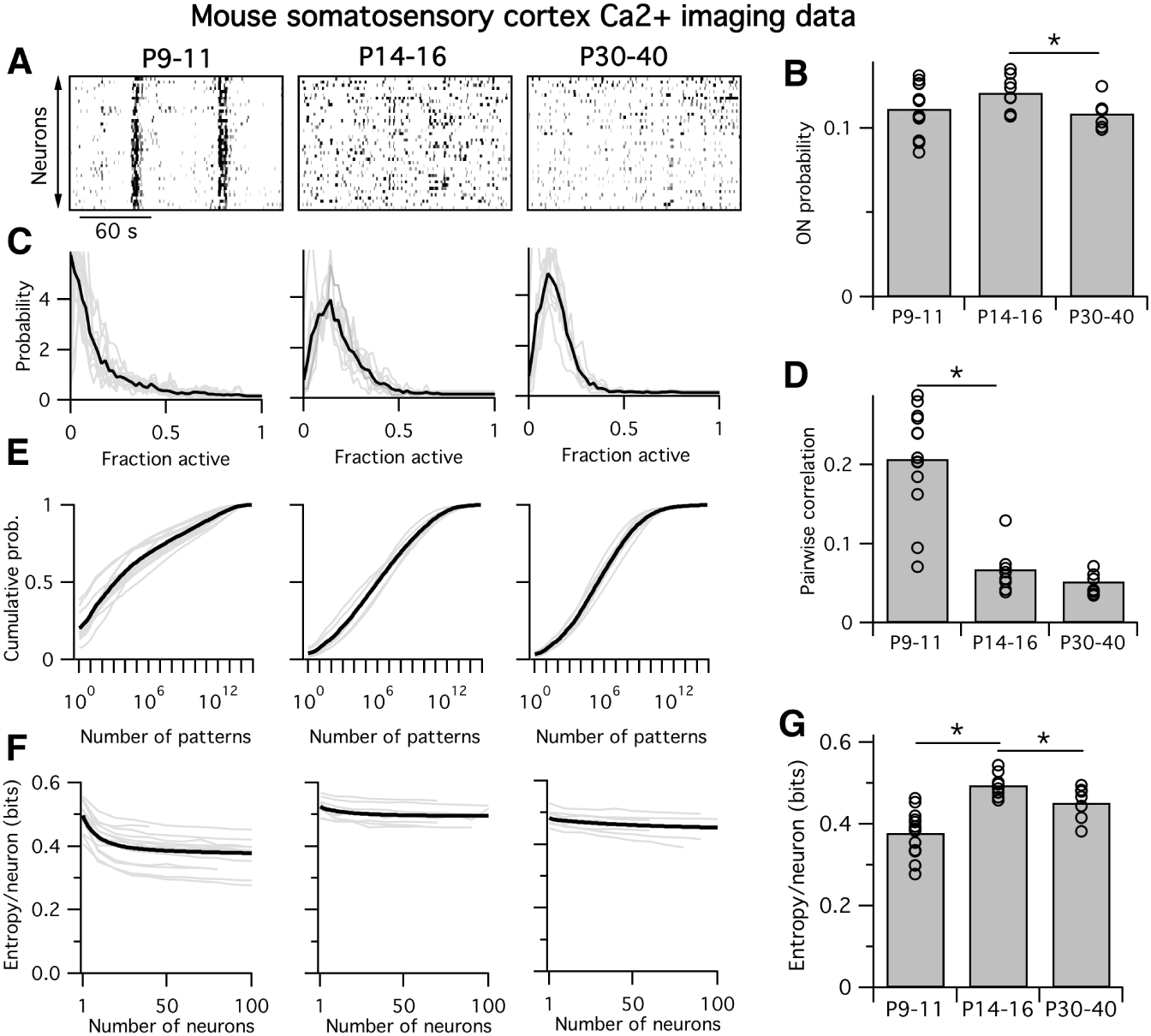
Entropy of neural populations in mouse somatosensory cortex increases then decreases during development. **A:** Example Ca^2+^ imaging movie from mice ages P9–11 (left), P14–16 (center), and P30–40 (right). **B:** Mean ON probability of neurons by group. Each circle corresponds to the mean across all neurons recorded in a single animal, bars represent group means. **C:** Probability density of the fraction of active neurons, for sets of 50 neurons. Light gray traces are distributions from single animals, heavy black traces are group means. **D:** Mean pairwise correlation between neurons in each group. **E:** Cumulative distribution of pattern probabilities for each group, for sets of 50 neurons. Note log scale on x-axes. **F:** Entropy per neuron as a function of the number of neurons analyzed. **G:** Estimate of mean entropy per neuron for 100 neurons.

We performed the analysis on data from mice at three developmental age points: P9–11 (n=13), P14–16 (n=8) and P30–40 (n=7). These correspond to timepoints just before (P9–P11) and after (P14–P16) the critical period for cortical plasticity, and mature stage post-weaning (P30–P40). Entropy is determined by two main properties of the neural population activity: the activity levels of the neurons and their correlations. We found than mean ON probability increased between ages P9–P11 and P14–16 (p=0.0016), then decreased again at age P30–40 (p=0.0024). As previously observed (Rochefort et al., 2009; Golshani et al., 2009; Gonçalves et al., 2013), mean pairwise correlations decreased across development (p<0.001, P9–P11 vs P14–P16) (Figure 8D) so that as animals aged there were fewer synchronous events when many neurons were active together (Figure 8A,C).

What do these statistics predict for the distribution of activity patterns exhibited by neural circuits? Interestingly, activity levels and correlations are expected to have opposite effects on entropy: in the sparse firing regime, any increase ON probability should increase the entropy by increasing the typical number of activity patterns due to combinatorics, while an increase in correlations should decrease the entropy because groups of neurons will tend to be either all ON or all OFF together.

When we quantified the entropy of the pattern probability distributions, we found a non-monotonic trajectory across development (Figure 8F–G). For 100-neuron populations, in young animals at P9–P11 we found a low group mean entropy of ∼ 0.38 bits/neuron (c.i. [0.347 : 0.406]), followed by an increase at P14–P16 (p<0.001) to ∼ 0.49 bits/neuron (c.i. [0.478 : 0.517]), and then a decrease in adulthood P30-P40 (p=0.036) to ∼ 0.45 bits/neuron (c.i. [0.418 : 0.476]). Although these shifts in dimensionality were subtle as estimated by entropy, they correspond to exponentially large shifts in pattern number. For example, 100-neuron populations in P14–P16 animals showed an average of 5.6 × 10^10^ patterns while 100-neuron populations in P30–P40 animals showed an 8-fold fewer number of ∼ 7.1 × 10^9^ typical patterns (data not shown). One interpretation of these findings is that young animals compress their neural representations of stimuli into a small ‘dictionary’ of activity patterns, then expand their representations into a larger dictionary at P14–P16, before again reducing the coding space again in adulthood, P30–P40.

Is the shift in cortical neural population entropy across development due to changes in firing rates, correlations, or both? We assessed this by fitting two control models to the same Ca^2+^ imaging data: the independent neuron model and the homogeneous population model (Figure 9). The independent neuron model captures changes in neural firing rates across development, including the heterogeneity in firing rates across the population, but inherently assumes that all correlations are fixed at zero. Although the independent model predicted a significant decrease in entropy between P14–16 and P30–40 (p=0.014) similar to the population tracking model, it did not detect an increase in entropy from P9–11 to P14–16 (p=0.13) (Figure 9B, left).

**Fig. 9:**
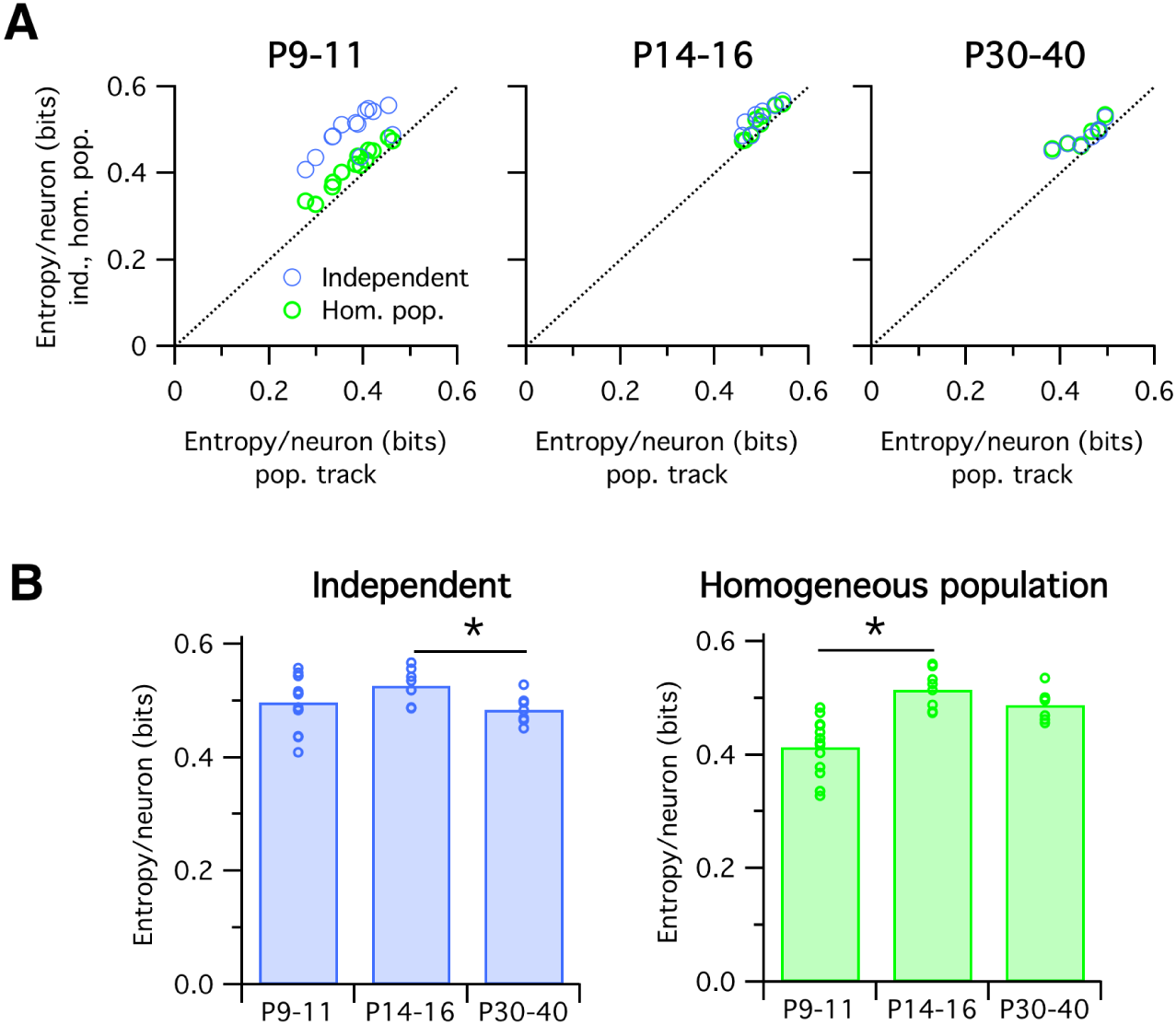
Mouse somatosensory cortex entropy trajectories are not captured by either the independent or homogeneous population models. **A:** Entropy per neuron estimated from the independent (blue circles) or homogeneous population (green) models against the same quantity estimated from the population tracking model, for data from mice of three age groups (left, center and right plots). Each circle indicates the joint entropy estimated for 100 neuron population recording from a single animal. Note that the independent and homogeneous population models always estimate greater entropy values than the population tracking model. **B:** Same data as panel A, plotted to compare to previous Figure 8G. Note that neither the independent (blue, left) nor homogeneous population (green, right) models predict the inverted-U shaped trajectory uncovered by the population tracking model (Figure 8G).

The homogeneous population model captures a different set of statistics. By matching the population synchrony distribution, it fits both the mean neuron firing rates and mean pairwise correlations. However, it also assumes that all neurons have identical firing rates and identical correlations, hence it does not capture any of the population heterogeneity that the independent neuron model does. In contrast to the independent model, the homogeneous population model did predict the increase in entropy from P9–11 to P14–16 (p=0.002), but did not detect a decrease in entropy from P14–16 to P30–40 (p=0.24).

Importantly, the independent and homogeneous population models always estimate greater entropy values than the population tracking model. This is to be expected since the population tracking model matches the key statistics of both control models together, and so cannot have a greater entropy than either. Together, these results demonstrate that the population tracking model can detect shifts in population entropy that could not be detected from either independent or homogeneous population models alone.

## 3 Discussion

Here we introduced a novel statistical model for neural population data. The model works by matching two features of the data: first, the probability distribution for the number of neurons synchronously active, and second, the conditional probability that each individual neuron is ON given the total number of active neurons in the population. The former set of parameters are informative about the general statistics of the population activity: the average firing rates and the level of synchrony. The latter set of parameters tell us more about the heterogeneity within the population: some neurons tend to follow the activity of their neighbors, while others tend to act independently. These two types of cells recently have been called ‘choristers’ and ‘soloists’, respectively (Okun et al., 2015).

Compared to existing alternatives (Table 1), the model we propose has several strengths: 1) it is rich enough to accurately predict pattern probabilities, even for large neural populations; 2) its parameters are computationally cheap to fit for large *N*; 3) the parameter estimates converge within an experimentally reasonable number of data timepoints, 4) sampling from the model is straightforward, with no correlation between consecutive samples; 5) it is readily normalizable to directly obtain pattern probabilities; 6) the model’s form permits a computationally tractable low-parameter approximation of the entire pattern probability distribution.

These strengths make the model appealing for certain neurobiological problems. However, since a pattern probability distribution can only be fully specified by 2^*N*^ numbers — so including correlation at all orders — whereas our model has only *N*^2^ parameters, it must naturally also have some shortcomings. The main weaknesses are: 1) since the population synchrony distribution becomes more informative with greater *N*, our model will in most cases be outperformed by alternatives for small *N*; 2) although ourmodel captures themean pairwise correlation across the population, it does not account for the full pairwise correlation structure (Figure 2C, center); 3) since the model considers only spatial correlations, temporal correlations are unaccounted for (Figure 2C, right); 4) The model parameters are not readily interpretable in a biological sense, unlike the pairwise couplings of the maximum entropy models (Schneidman et al., 2006), or the stimulus filters in Generalized Linear Models (Pillow et al., 2008); 5) unlike classic maximum entropy models, ours carries no notion of an energy landscape and so does not imply a natural dynamics across the state space (Tkacik et al., 2014).

We demonstrated the utility of the population tracking model by applying it to two neurobiological problems. First, we found that the population tracking model allowed fast prediction of visual stimuli by decoding neural population data from macaque primary visual cortex (Figure 7). A simple but widely used alternative model that assumes independent neurons achieved 50% decoding accuracy around 20 ms after performance rose above chance levels. In contrast,. the population tracking model reached 50% accuracy only ∼ 14 ms after exceeding chance levels. Since we binned time in 10 ms intervals, this implies that the population tracking model was correct more often than not given neural population data from less than two timepoints, on average. What does this finding imply for brain function? The actual decoding algorithm we used for this task, Maximum Likelihood, is not neurobiologically plausible. However, the fact that the population tracking model worked so well implies two things about cortical visual processing. First, sufficient information is present in the spiking patterns of these neural populations to perform stimulus discrimination very quickly after the stimulus response onset. Previous studies found that good decoding performance for similar tasks was typically achieved at least 80–100 ms following stimulus onset (Chen et al., 2008; Berens et al., 2012), whereas the population tracking model took only ∼ 65 ms. However, direct comparisons with these previous studies are problematic: for example, on the one hand Berens et al. (2012) examined only 20 units while we considered groups up to *N* = 100, but on the other hand Berens et al. (2012) considered only a binary classification task whereas we considered the more difficult task of decoding a single stimulus orientation from all eight possibilities. Further work is needed to resolve these issues. Second, the improved performance of the population tracking model over the independent model implies that it may be beneficial for the brain to explicitly represent the number of neurons simultaneously active in the local circuit. Indeed this seems like a natural computation for single neurons to perform as they sum the synaptic inputs from their neighboring neurons. Our finding implies that this summed value itself carries additional information about the stimulus beyond that present in the list of identities of active neurons. Whether and how the brain uses this information remain questions for future study.

Our second application of the population tracking model was to look for changes in the distribution of neural pattern probabilities in mouse somatosensory cortex across development (Figure 8). We found a surprising non-monotonic trajectory across development. Initially at P9–11 the entropy of population activity is low, due to large synchronous events in the population. The correlations decrease dramatically at around P12 (Golshani et al., 2009; Rochefort et al., 2009), so that at P14–16 activity is relatively desynchronized, leading to an increase in population entropy. However, we then found a reduction in firing rates from P14–16 to P30–40 that corresponded to a decrease in entropy, despite no large change in correlations. These findings uncover a subtle and unexplained developmental trajectory for mouse somatosensory cortex that warrants detailed further study. Importantly, this nonmonotonic development curve would not have been detectable by examining either firing rates or correlations in isolation (Figure 9).

The population tracking model we propose is similar in spirit to a recently proposed alternative, the population coupling model (Okun et al., 2012, 2015; Schölvinck et al., 2015). These authors developed a model of neural population data with *3N* parameters: *N* specifying the firing rates of each neuron, another *N* specifying the population rate distribution, and a final *N* specifying the linear coupling of each individual neuron with the population rate. Okun et al. (2015) fit this model to data from mouse, rat, and primate cortex and found that neighboring neurons showed diverse couplings to the population rate, that this coupling was invariant to stimulus conditions, and that the degree of a neuron’s population coupling was reflected in the number of synaptic inputs it received from its neighbors. These results show that the population rate contains valuable statistical information that can help constrain models of neural population dynamics. Despite these notable advances, the population coupling model of Okun et al. also suffers from several shortcomings that our model does not: first, it offers no way to write down either the probability of a single neural activity pattern or the relative probabilities of two activity patterns in terms of the model’s parameters. Second, for large neural populations there is no way to estimate functions of the entire pattern probability distribution, such as the Shannon entropy or the Kullback-Leibler divergence. Third, generating samples from the model involves a computationally expensive iterative procedure, and the probability distribution across possible samples is not fully determined by the model parameters, but depends also on the experimenter’s choice of sampling algorithm. Finally, the model assumes a linear relationship between each individual neuron’s firing rate and the population rate. Although parsimonious, this linear model may be insufficiently flexible to capture the true relationship. Also a linear model must break down at some point: a neuron cannot fire at rates less than zero Hertz or at rates higher than its maximal firing frequency. For all of these reasons, we suggest that the model we propose may be applicable to a wider range of neurobiological problems than the population coupling model.

In what scenarios will the population tracking model do best and worst in? Intuitively, the model will do best when the true pattern probability distribution, which in principle could take any arbitrary shape in its 2^*N*^-dimensional space, is nearby to the family of probability distributions that are attainable from the population tracking model, which has only *N*^2^ degrees of freedom. A rigorous mathematical understanding of the neural activity regimes that could be well-matched by the population tracking model remains a goal for future studies. Nevertheless, we can hazard an answer to this question based on the form of the model. Given that the population tracking model assumes that all individual neurons are coupled only via a single global population rate variable *K*, it will be unlikely that the model can well capture any correlations within or between any specific subgroups present in the data. Presumably the degree of error that this introduces will increase with increasing heterogeneity in correlation structure, especially if the neural population is highly modular. Indeed we found that the entropy estimated for heterogeneous DG model samples was less accurate than the case where DG model parameters were more homogeneous (compare Figure 4D, left with Figure 6C). We do note however that the population tracking model can capture some of the pairwise correlation structure beyond the means, as observed in Figure 2C and Appendix Figure 1. This may be due to the fact that the model captures the heterogeneity in firing rates, which can affect pairwise correlations (de la Rocha et al., 2007). Overall, we suggest that the primary benefit of the population tracking model may not be that it is the most accurate of all available models, but that it preserves its accuracy and tractability for large *N* datasets.

What type of new neurobiological research questions can we ask with the population tracking model? We introduced a method for calculating the divergence between the model fits to two sets of neural population activity data. This measure should be useful for experiments where the same neurons are recorded in two or more different conditions, such as comparing the statistics of spontaneous activity with that evoked by stimuli (Figure 5), or the effects of an acute pharmacological or optogenetic stimulation on neural circuit activity. In contrast, if experiments involve comparing neural population activity from different animals, such as genetically distinct animals or at different timepoints in development, one can still perform quantitative comparisons of the activity statistics at a grouped population level (Figure 8).

The most direct usage of our model may however be to provide limits and constraints on future theoretical models of neural population coding. The Shannon entropy is a particularly useful measure because it provides an upper bound on the information that the neural population can represent. We conjecture, but have not proven, that our model is maximum entropy given the parameters. Adding temporal correlations, which real neurons show but are not included in the population tracking model, can only further reduce the population entropy. Hence, assuming that enough data are available for the model parameter fits to converge, the entropy estimate from the population tracking model gives a hard upper bound on the coding capacity of a circuit. Any feasible model for neural processing in a given brain region must obey these limits.

## Acknowledgements

We thank Conor Houghton, Timothy O’Leary, Hannes Saal, and Alex Williams for comments on earlier versions of the manuscript. This study was supported by funding from FRAXA Research Foundation, Howard Hughes Medical Institute, Sloan-Swartz Foundation, the Dana Foundation, and the NIH (NICHD R01HD054453 and NINDS RC1NS068093). The macaque recordings from the laboratory of Adam Kohn were funded by NIH grant EY016774.

## Appendix

### Macaque electrophysiological recording

All macaque electrophysiology data were previously published (Zandvakili and Kohn, 2015) and kindly shared by A. Kohn. Full details of experimental procedures and raw data processing steps are available in Zandvakili and Kohn (2015).

### Mouse in vivo calcium imaging recording

All Ca^2+^ imaging data were previously published (Gonçalves et al., 2013). Briefly, data were collected from male and female C57Bl/6 wild-type mice at P9–40. Mice were anesthetized with isoflurane, and a cranial window was fitted over primary somatosensory cortex by stereotaxic coordinates. Mice were then transferred to a two-photon microscope and headfixed to the stage while still under isoflurane anesthesia. 2–4 injections of the Ca^2+^ sensitive Oregon-Green BAPTA-1 (OGB) dye and sulforhodamine-101 (to visualize astrocytes) were injected 200 μm below the dura. Calcium imaging was performed using a Ti-Sapphire Chameleon Ultra II laser (Coherent) tuned to 800 nm. Imaging in unanesthetized mice began within 30-60 mins of stopping the flow of isoflurane after the last OGB injection. Images were acquired using ScanImage software (Pologruto et al., 2003) written in MATLAB (MathWorks). Whole-field images were collected using a 20× 0.95 NA objective (Olympus) at an acquisition speed of 3.9 Hz (512 × 128 pixels).

Several 3-minute movies were concatenated and brief segments of motion artifacts were removed (always <10 s total). Data were corrected for *x-y* drift. Cell contours were automatically detected and the average Δ*F/F* signal of each cell body was calculated at each time point. Each Δ*F/F* trace was low-pass filtered using a Butterworth filter (coefficient of 0.16) and deconvolved with a 2 s single-exponential kernel (Yaksi and Friedrich, 2006). To remove baseline noise, the standard deviation of all points below zero in each deconvolved trace was calculated, multiplied by two, and set as the positive threshold level below which all points in the deconvolved trace were set to zero. Estimated firing rates of the neurons, *r_i_*(*t*), were then obtained by multiplying the deconvolved trace by a factor of 78.4, which was previously derived empirically from cell-attached recordings *in vivo* (Golshani et al., 2009).

### Data analysis methods

All data analysis and calculations were done using MATLAB (The Mathworks).

#### Statistical tests

To avoid parametric assumptions, all statistical tests were done using standard bootstrapping methods with custom-written MATLAB scripts. For example when assessing the observed difference between two group means Δ_*μobs*_ we performed the following procedure to calculate a p-value. First we pool the data points from the two groups to create a null set *S_null_*. We then construct two hypothetical groups of samples *S*_1_ and S_2_ from this by randomly drawing *n*_1_ and *n*_2_ samples with replacement from *S_null_*, where *n*_1_ and *n*_2_ are the number of data points in the original groups 1 and 2 respectively. We take the mean of both hypothetical sets *μ*_1_ and *μ*_2_ and calculate their difference Δ*μ*_*null*_ = *μ*_1_ − *μ*_2_. We then repeat the entire procedure 10^7^ times to build up a histogram of Δ*μ*_*null*_. This distribution is always centered at zero. After normalizing, this can be interpreted as the probability distribution Pr(Δ*μ*_*null*_) for observing a group mean difference of Δ*μ*_*null*_ purely by chance if the data were actually sampled from the same null distribution. Then the final p-value for the probability of finding a group difference of at least Δ*μ*_*obs*_ in either direction is given by

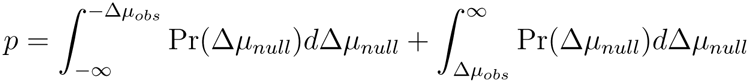

Any data that varied over multiple orders of magnitude (e.g. the number of patterns observed) was log-transformed before comparing group means.

#### Conversion from firing rate to ON/OFF probabilities for Ca^2+^ imaging data

For the Ca^2+^ imaging data, we began with estimated firing rate time series *r_i_*(*t*) for each neuron i recorded as part of a population of *N* neurons. For later parts of the analysis we needed to convert these firing rates to binary ON/OFF values. This conversion involves a choice. One option would be to simply threshold the data, but this would throw away information about the magnitude of the firing rate. We instead take a probabilistic approach where rather than deciding definitively whether a given neuron was ON or OFF in a given time bin, we calculate the probability that the neuron was ON or OFF by assuming that neurons fire action potentials according to an inhomogeneous Poisson process with rate *r_i_*(*t*). The mean number of spikes λ_*i*_(*t*) expected in a time bin of width Δ*t* is λ_*i*_(*t*) = *r_i_*(*t*) × Δ*t*. We choose Δ*t* = 1 second. Under the Poisson model the actual number of spikes m in a particular time bin is a random variable that follows the Poisson distribution 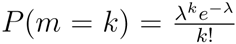. We will consider a neuron active (ON) if it is firing one or more spikes in a given time bin. Hence the probability that a neuron is ON is *p_on_*(*t*) = 1 – P(*m* = 0) = 1 – *e*^λ^. This approach has two advantages over thresholding: 1) it preserves some information about the magnitude of firing rates, and 2) it acts to regularize the probability distribution for the number of neurons active by essentially smoothing nearby values together.

#### Entropy estimation for large numbers of neurons for Ca^2+^ imaging data

The entropy/neuron generally decreased slightly with the number of neurons considered as result of the population correlations (see Figure 8F in main text), so we needed to control for neural population size when comparing data from different experimental groups. On the one hand we would like to study as large a number of neurons as possible, because we expect the effects of collective network dynamics to be stronger for large population sizes and this may be the regime where differences between the groups emerge. On the other hand our recording methods allowed us to sample only typically around ∼ 100 neurons at a time, and as few as 40 neurons in some animals. Hence we proceeded by first estimating the entropy/neuron in each animal by calculating the entropy of random subsets of neurons of varying size from 10 to 100 (if possible) in steps of 10. For each population size we sampled a large number of independent subsets, calculated the entropy of each. Finally for each dataset we fit a simple decaying exponential function to the entropy/neuron as a function of the number of neurons: 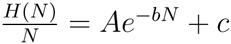, and used this fit to estimate *H/N* for 100 neurons.

**Appendix Figure 1:**
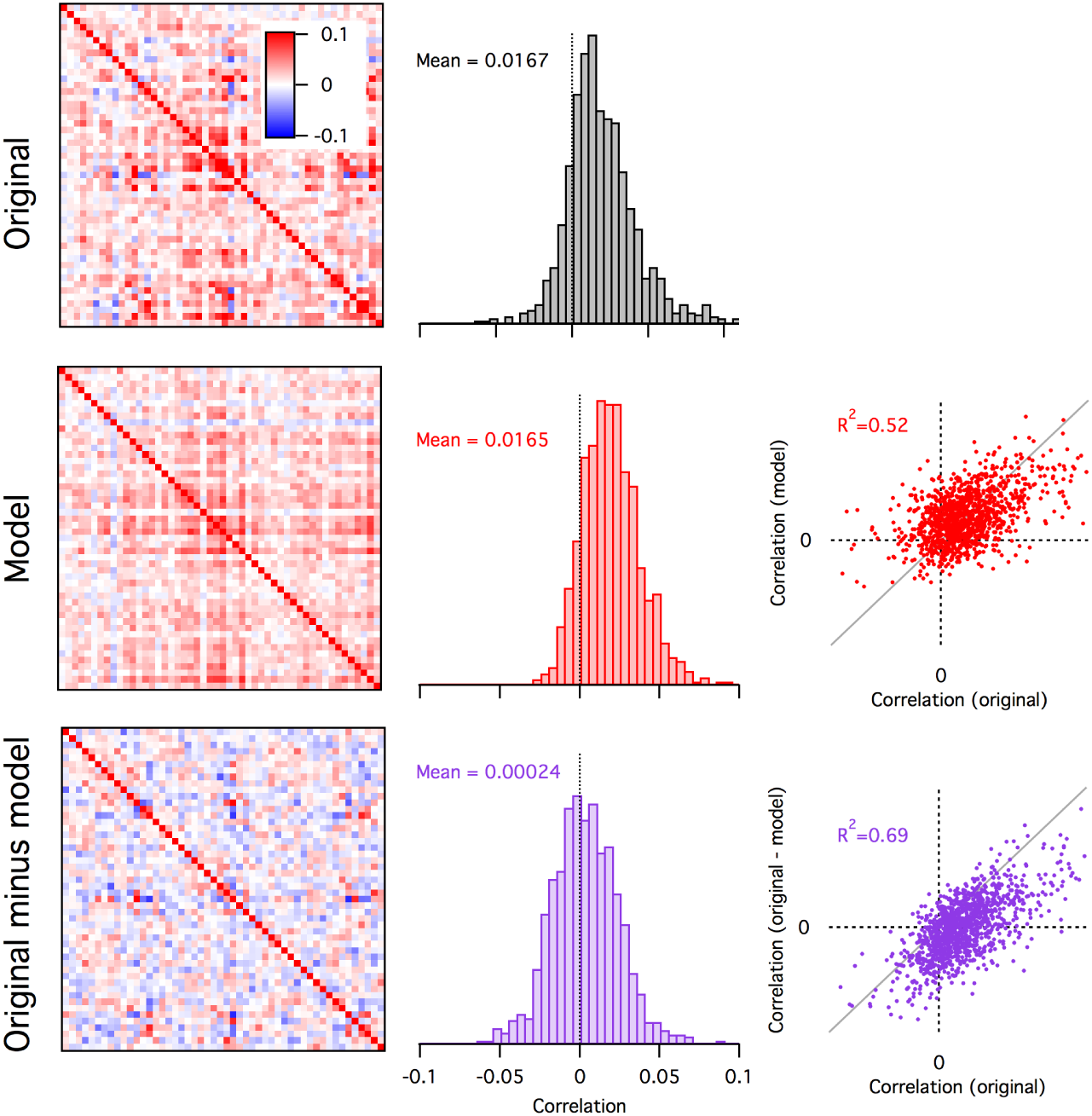
The population tracking model partially recapitulates the pairwise correlation structure of the original data. Left column are the pairwise correlation matrices from the example data shown in Figure 2 (top), for samples drawn from the population tracking model fit to these data (center), and the residual pairwise correlations in the data after subtracting the covariance accounted for by the population tracking model and renormalizing (bottom). Center column are histograms of the pairwise correlations from each matrix in the left column. The scatter plots in the right column show the individual pairwise correlations of the model (red) and the data minus the model (purple) against the pairwise correlations in the original data. Note that the model almost exactly captures the mean pairwise correlation of the original data, and part of the remaining structure (R^2^ = 0.52).

## References

Amari, S.-I., Nakahara, H., Wu, S., and Sakai, Y. (2003). Synchronous firing and higher-order interactions in neuron pool. Neural computation.

Archer, E. W., Park, I. M., and Pillow, J. W. (2013). Bayesian entropy estimation for binary spike train data using parametric prior knowledge. Advances in neural information….

Averbeck, B. B., Latham, P. E., and Pouget, A. (2006). Neural correlations, population coding and computation. Nature reviews Neuroscience.

Berens, P., Ecker, A. S., Cotton, R. J., Ma, W. J., Bethge, M., and Tolias, A. S. (2012). A fast and simple population code for orientation in primate V1. The Journal of neuroscience : the official journal of the Society for Neuroscience.

Berkes, P., Orbán, G., Lengyel, M., and Fiser, J. (2011). Spontaneous cortical activity reveals hallmarks of an optimal internal model of the environment. Science.

Berry, M. J. II, Tkacik, G., Dubuis, J., Marre, O., and da Silveira, R. A. (2013). A simple method for estimating the entropy of neural activity. Journal of Statistical Mechanics: Theory and Experiment.

Broderick, T., Dudik, M., Tkacik, G., Schapire, R. E., and Bialek, W. (2007). Faster solutions of the inverse pairwise Ising problem. arXiv.org.

Buzsáki, G. and Mizuseki, K. (2014). The log-dynamic brain: how skewed distributions affect network operations. Nature reviews Neuroscience.

Chen, Y., Geisler, W. S., and Seidemann, E. (2006). Optimal decoding of correlated neural population responses in the primate visual cortex. Nature neuroscience.

Chen, Y., Geisler, W. S., and Seidemann, E. (2008). Optimal temporal decoding of neural population responses in a reaction-time visual detection task. Journal of neurophysiology.

Churchland, P. S. and Sejnowski, T. J. (1994). The Computational Brain. Mit Press.

Cohen, M. R. and Kohn, A. (2011). Measuring and interpreting neuronal correlations. Nature neuroscience.

Cover, T. M. and Thomas, J. A. (2006). Elements of Information Theory. Wiley-Interscience.

Cui, Y., Liu, L. D., McFarland, J. M., Pack, C. C., and Butts, D. A. (2016). Inferring Cortical Variability from Local Field Potentials. The Journal of neuroscience : the official journal of the Society for Neuroscience.

Cunningham, J. P. and Yu, B. M. (2014). Dimensionality reduction for large-scale neural recordings. Nature neuroscience.

de la Rocha, J., Doiron, B., Shea-Brown, E., Josić, K., and Reyes, A. (2007). Correlation between neural spike trains increases with firing rate. Nature.

Ganmor, E., Segev, R., and Schneidman, E. (2011). Sparse low-order interaction network underlies a highly correlated and learnable neural population code.

Gerstein, G. L. and Perkel, D. H. (1969). Simultaneously recorded trains of action potentials: analysis and functional interpretation. Science.

Gerstein, G. L. and Perkel, D. H. (1972). Mutual temporal relationships among neuronal spike trains. Statistical techniques for display and analysis. Biophysical journal.

Golshani, P., Gonçalves, J. T., Khoshkhoo, S., Mostany, R., Smirnakis, S., and Portera-Cailliau, C. (2009). Internally mediated developmental desynchronization of neocortical network activity. The Journal of neuroscience : the official journal of the Society for Neuroscience.

Gonçalves, J. T., Anstey, J. E., Golshani, P., and Portera-Cailliau, C. (2013). Circuit level defects in the developing neocortex of Fragile X mice. Nature neuroscience.

Köster, U., Sohl-Dickstein, J., Gray, C. M., and Olshausen, B. A. (2014). Modeling higher-order correlations within cortical microcolumns. PLoS computational biology.

Macke, J. H., Berens, P., Ecker, A. S., Tolias, A. S., and Bethge, M. (2009). Generating spike trains with specified correlation coefficients. Neural computation.

Macke, J. H., Murray, I., and Latham, P. E. (2011a). How biased are maximum entropy models? Advances in neural information….

Macke, J. H., Opper, M., and Bethge, M. (2011b). Common input explains higher-order correlations and entropy in a simple model of neural population activity. Physical review letters.

Marre, O., El Boustani, S., Frégnac, Y., and Destexhe, A. (2009). Prediction of spatiotemporal patterns of neural activity from pairwise correlations. Physical review letters.

Nasser, H., Marre, O., and Cessac, B. (2013). Spatio-temporal spike train analysis for large scale networks using the maximum entropy principle and Monte Carlo method. Journal of Statistical Mechanics: Theory and Experiment.

Ohiorhenuan, I. E., Mechler, F., Purpura, K. P., Schmid, A. M., Hu, Q., and Victor, J. D. (2010). Sparse coding and high-order correlations in fine-scale cortical networks. Nature.

Okun, M., Steinmetz, N. A., Cossell, L., Iacaruso, M. F., Ko, H., Bartho, P., Moore, T., Hofer, S. B., Mrsic-Flogel, T. D., Carandini, M., and Harris, K. D. (2015). Diverse coupling of neurons to populations in sensory cortex. Nature.

Okun, M., Yger, P., Marguet, S. L., Gerard-Mercier, F., Benucci, A., Katzner, S., Busse, L., Carandini, M., and Harris, K. D. (2012). Population rate dynamics and multineuron firing patterns in sensory cortex. The Journal of neuroscience : the official journal of the Society for Neuroscience.

Park, I. M., Archer, E. W., Latimer, K., and Pillow, J. W. (2013). Universal models for binary spike patterns using centered Dirichlet processes. Advances in neural….

Perkel, D. H., Gerstein, G. L., and Moore, G. P. (1967). Neuronal spike trains and stochastic point processes. II. Simultaneous spike trains. Biophysical journal.

Pillow, J. W., Shlens, J., Paninski, L., Sher, A., Litke, A. M., Chichilnisky, E. J., and Simoncelli, E. P. (2008). Spatio-temporal correlations and visual signalling in a complete neuronal population. Nature.

Pnevmatikakis, E. A., Soudry, D., Gao, Y., Machado, T. A., Merel, J., Pfau, D., Reardon, T., Mu, Y., Lacefield, C., Yang, W., Ahrens, M., Bruno, R., Jessell, T. M., Peterka, D. S., Yuste, R., and Paninski, L. (2016). Simultaneous Denoising, Deconvolution, and Demixing of Calcium Imaging Data. Neuron.

Pologruto, T. A., Sabatini, B. L., and Svoboda, K. (2003). ScanImage: flexible software for operating laser scanning microscopes. Biomedical engineering online.

Quiroga, R. Q. (2012). Spike sorting. Current biology : CB.

Rahmati, V., Kirmse, K., Marković, D., Holthoff, K., and Kiebel, S. J. (2016). Inferring Neuronal Dynamics from Calcium Imaging Data Using Biophysical Models and Bayesian Inference. PLoS computational biology.

Rochefort, N. L., Garaschuk, O., Milos, R.-I., Narushima, M., Marandi, N., Pichler, B., Kovalchuk, Y., and Konnerth, A. (2009). Sparsification of neuronal activity in the visual cortex at eye-opening.

Roudi, Y., Nirenberg, S., and Latham, P. E. (2009). Pairwise maximum entropy models for studying large biological systems: when they can work and when they can’t. PLoS computational biology.

Schaub, M. T. and Schultz, S. R. (2012). The Ising decoder: reading out the activity of large neural ensembles. Journal of computational neuroscience.

Schneidman, E., Berry, M. J., Segev, R., and Bialek, W. (2006). Weak pairwise correlations imply strongly correlated network states in a neural population. Nature.

Schölvinck, M. L., Saleem, A. B., Benucci, A., Harris, K. D., and Carandini, M. (2015). Cortical state determines global variability and correlations in visual cortex. The Journal of neuroscience : the official journal of the Society for Neuroscience.

Shlens, J., Field, G. D., Gauthier, J. L., Grivich, M. I., Petrusca, D., Sher, A., Litke, A. M., and Chichilnisky, E. J. (2006). The structure of multi-neuron firing patterns in primate retina. The Journal of neuroscience : the official journal of the Society for Neuroscience.

Singer, W. (1999). Neuronal Synchrony: A Versatile Code for the Definition of Relations? Neuron.

Stevenson, I. H. and Kording, K. P. (2011). How advances in neural recording affect data analysis. Nature neuroscience.

Tang, A., Jackson, D., Hobbs, J., Chen, W., Smith, J. L., Patel, H., Prieto, A., Petrusca, D., Grivich, M. I., Sher, A., Hottowy, P., Dabrowski, W., Litke, A. M., and Beggs, J. M. (2008). A maximum entropy model applied to spatial and temporal correlations from cortical networks in vitro. The Journal of neuroscience : the official journal of the Society for Neuroscience.

Tkacik, G., Marre, O., Amodei, D., Schneidman, E., Bialek, W., and Berry, M. J. (2014). Searching for collective behavior in a large network of sensory neurons. PLoS computational biology.

Tkacik, G., Marre, O., Mora, T., Amodei, D., Berry II, M. J., and Bialek, W. (2013). The simplest maximum entropy model for collective behavior in a neural network. Journal of Statistical Mechanics: Theory and Experiment.

Yaksi, E. and Friedrich, R. W. (2006). Reconstruction of firing rate changes across neuronal populations by temporally deconvolved Ca2+ imaging. Nature methods.

Yeh, F.-C., Tang, A., Hobbs, J., Hottowy, P., Dabrowski, W., Sher, A., Litke, A., and Beggs, J. (2010). Maximum Entropy Approaches to Living Neural Networks. Entropy.

Yu, S., Huang, D., Singer, W., and Nikolić, D. (2008). A small world of neuronal synchrony. Cereb Cortex.

Yu, S., Yang, H., Nakahara, H., Santos, G. S., Nikolić, D., and Plenz, D. (2011). Higher-order interactions characterized in cortical activity. The Journal of neuroscience : the official journal of the Society for Neuroscience.

Zandvakili, A. and Kohn, A. (2015). Coordinated Neuronal Activity Enhances Corticocortical Communication. Neuron.

Zohary, E., Shadlen, M. N., and Newsome, W. T. (1994). Correlated neuronal discharge rate and its implications for psychophysical performance. Nature.

